# Beyond Aging, Sex and Insomnia Disorder Shape NREM Brain Oscillations

**DOI:** 10.64898/2026.03.17.712450

**Authors:** Nyissa A. Walsh, Aurore A. Perrault, Nathan E. Cross, Antonia Maltezos, Emma-Maria Phillips, Loïc Barbaux, Oren Weiner, Claire Dyment, Florence Borgetto, Jean-Phillipe Gouin, Thien Thanh Dang-Vu

**Author notes:** Corresponding authors, Concordia University, 7141 Rue Sherbrooke O, H4B 1R6, Montreal, QC, Canada, Fax: N/A.

## Abstract

**Objectives:** Chronic insomnia (INS) is particularly prevalent in older adults and females. Sex-and age-related differences in neurophysiological markers of sleep quality (sleep spindles and slow-wave activity [SWA]) may underlie differential vulnerability to INS. This study investigated the effects of sex and insomnia on spindle and SWA beyond aging, to better understand the mechanistic differences contributing to the higher prevalence of INS in females.

**Methods:** After a habituation night, one night of sleep assessed with polysomnography was analyzed in 222 adults (aged 18-82) including 119 INS (71% female) and 103 healthy sleepers (HS; 61% female). Spindle density, slow oscillation (SO) density, relative sigma power and SWA were derived during NREM sleep. Age, group, sex, and group-by-sex interactions were examined, with age as a covariate.

**Results:** Age, insomnia, and sex each contributed uniquely to NREM oscillatory activity. INS primarily reduced spindle and SO density, while sex accounted for differences in SWA. While SWA was higher in females overall, sex differences were not significant within the INS or HS groups. Female INS reported highest rates of insomnia severity as well as lower sigma power than males in the INS group. Spindle and SO density deficits were also present in female INS relative to female HS, as well as male INS relative to male HS.

**Conclusions:** The combination of reduced sigma power in females with insomnia relative to their male counterparts, as well as less spindle and SO density compared to female healthy sleepers may contribute to greater insomnia severity in females.

**Statement of Significance:** Insomnia is a growing public health concern that is more commonly reported in females, yet the neural mechanisms underlying this sex difference remain poorly understood. Our findings suggest that specific markers of sleep quality are disproportionately disrupted in females with insomnia, potentially contributing to greater vulnerability and symptom severity. These results provide new insight into how sex influences the neurophysiology of insomnia disorder and identify oscillatory markers that could serve as targets for personalized interventions. Future research should investigate whether these alterations represent persistent dysfunction or reversible changes, which could advance understanding of the biological basis of insomnia and inform strategies to improve sleep health in at-risk populations.

## Introduction

Chronic insomnia is a growing public health concern, affecting approximately 16% of Canadians, and individuals worldwide^1,2^. Chronic insomnia is defined by significant daytime impairments and reduced quality of life^3–5^ as a result of persistent difficulty initiating or maintaining sleep, at least three nights a week, for at least three months despite adequate opportunity for rest.

Although chronic insomnia diagnosis is based on self-reported complaints, altered objective sleep measures can also be commonly observed within this population. Polysomnography (PSG) sleep studies show that, relative to healthy sleepers, some individuals with chronic insomnia exhibit prolonged sleep onset latency, increased time in lighter stages of non-rapid eye movement (NREM) sleep (i.e., NREM1 and NREM2), frequent nocturnal awakenings, and reduced sleep efficiency^6–8^. Beyond conventional macro-architectural measures, NREM sleep is defined by specific neural oscillations that reflect the integrity and restorative properties of sleep at a neurophysiological level and may therefore provide a more sensitive marker of insomnia-related sleep disruption^9–11^. Sleep spindles are brief (0.5 to 3s) oscillating bursts of 11–16Hz sigma activity that can be observed frontally, and over central–parietal regions^12,13^. Slow wave activity (SWA) is composed of delta waves oscillating between 1-4Hz as well as high-voltage biphasic waves called slow oscillations (SO; <1.25Hz)^14–16^. These rhythms, while mainly known for their implications in sleep-dependent memory processes^17,18^, also play a central role in maintaining sleep continuity and depth^19–21^. Reduced spindle activity is a reliable marker of disrupted sleep^12,22,23^, and some studies have shown that insomnia is also associated with reduced delta and SO power, increased fast-frequency EEG activity, and blunted slow-wave rebound following sleep deprivation, patterns consistent with hyperarousal and altered sleep homeostasis^9,16,24,25^.

Importantly, sleep quality and its underlying neurophysiological oscillations are shaped by key biological moderators, most notably sex and age. In healthy populations females generally exhibiting greater sleep spindle density, and higher absolute sigma and delta power compared to males^26^. However, findings across studies have been mixed and appear to vary depending on methodology, age, and analytic approach^26,27^. Contrasting to the female sleep advantage in some NREM brain oscillations, epidemiological studies consistently demonstrate that chronic insomnia is more prevalent in females than males, with females also showing a tendency to perceive and experience insomnia as more distressing and impairing^1,28^. Importantly, while sex differences in sleep oscillations are well-documented among healthy sleepers^26,29,30^, it remains unclear how these differences manifest in individuals with disturbed sleep, such as those with chronic insomnia.

In addition to sex, aging also influences sleep physiology, further shaping the quality and stability of sleep across the lifespan. Both SWA and sleep spindle characteristics decline with age due to structural and functional brain changes, including neuronal loss, neurobiological deterioration, reduced thalamocortical connectivity, diminished GABAergic signaling, and cortical thinning^31–33^. Natural age-related changes in sex hormones further modulate these processes. For instance, estradiol and progesterone enhance spindle activity via GABAergic mechanisms^34^, while SWA appears less sensitive to hormonal fluctuations, likely due to its regulation by sleep homeostasis^35–37^. Testosterone may also influence cortical structures involved in spindle and SWA, but its effects remain understudied^36,38,39^. These neurobiological and hormonal age-related changes contribute to shallower, more fragmented sleep and higher insomnia risk in older adults^33,40^. Moreover, sex disparities in insomnia prevalence become apparent during puberty and persist across the lifespan, with higher risks during reproductive events (i.e., menarche, pregnancy, post-partum, perimenopause)^41–44^. The highest incidence of insomnia occurs in midlife and older females^1,4^. During perimenopause and post-menopause (i.e., total cessation of menstruation), rates of insomnia increase likely due to hormonal changes such as estrogen decline, although comorbid health issues and psychosocial stressors may also contribute^45,46^. In healthy sleepers, age is associated with decreases in delta power, sigma power, and centro-parietal spindle density in both sexes^31,47,48^, but even with these declines, females tend to exhibit higher spindle density than males throughout the lifespan^31^.

To date, most research on sex differences in sleep neurophysiology has focused on healthy individuals across the lifespan, leaving gaps in our understanding of how the presence of insomnia may impact sleep micro-architecture. Given the critical role of sleep spindles, delta waves and SOs in sleep quality and maintenance, sex differences in brain oscillations may contribute to differential vulnerability for insomnia and shape its clinical presentation. The current study aims to investigate the effects of age, biological sex and presence of chronic insomnia on spindle and SWA by identifying unique contributions of each factor and their interactions in a large dataset of individuals with and without insomnia, accounting for the influence of age. We hypothesize that above and beyond age, individuals with chronic insomnia will exhibit reduced spindle and SO density, as well as reduced sigma power and SWA relative to healthy sleepers. However, when stratified by sex, we expect females with chronic insomnia to show lower spindle and SWA than males with chronic insomnia given the higher severity of complaints and neurobiological/hormonal changes in females across the lifespan.

## Method

### Participants

Data used for the current study were collected in the scope of six different projects (published^11,49–52^ or registered [ISRCTN13983243, NCT04024787, ISRCTN12645581] elsewhere) investigating sleep in individuals with chronic insomnia and healthy sleepers within the Sleep, Cognition, and Neuroimaging Laboratory in Montreal, Canada. All groups had a similar recruitment process, and participants were recruited through online and print advertisements, community postings, and physician referrals. Initial eligibility was assessed via a telephone screening, followed by a semi-structured in-person clinical interview, which most studies included the Structured Clinical Interview for DSM (SCID) to assess comorbid mental disorders. Eligibility criteria varied slightly across studies (e.g., age). Healthy sleepers were self-identified good sleepers, reported no sleep complaints, and had no history of sleep disorders. Participants with insomnia met diagnostic criteria for chronic insomnia disorder according to the *Diagnostic and Statistical Manual of Mental Disorders* (DSM-5^4^) and the *International Classification of Sleep Disorders, 3rd edition* (ICSD-3^3^). Criteria included self-reported difficulty initiating or maintaining sleep, or early morning awakenings occurring ≥3 nights per week for ≥3 months, with associated daytime impairment. General exclusion criteria for both groups included being outside the eligible age range (< 18 years), current neurological or psychiatric disorders other than anxiety or depression, medical conditions likely to affect sleep (e.g., epilepsy, multiple sclerosis, Parkinson’s disease, chronic pain, active cancer), untreated thyroid disorders, or major cardiovascular events (e.g., myocardial infarction, stroke). Other exclusion criteria included the presence of sleep disorders identified through moderate to severe sleep apnea (apnea-hypopnea index; AHI >5-15/h), and periodic limb movement (index >15/h). Participants were excluded for current use of medications known to affect sleep (e.g., hypnotics, antidepressants, and over-the-counter medication), inability to abstain from such medications for at least 1 week prior to the study, frequent alcohol (>10 drinks/week), cannabis or illicit drug use. Some studies included additional restrictions, such as differences in abstinence periods from prior medications or tobacco use. All participants provided written informed consent prior to participation. Procedures were approved by the Concordia University Human Research Ethics Committee and by the Comité d’Ethique de la Recherche of the Centre de Recherche de l’Institut de Gériatrie de Montréal (CRIUGM).

#### Insomnia Severity

Insomnia symptoms were assessed using the *Insomnia Severity Index* (ISI)^53^, a seven-item self-report instrument evaluating the nature, severity, and impact of insomnia over the preceding two weeks. Items assess difficulties with sleep onset, sleep maintenance, and early morning awakenings, as well as satisfaction with current sleep patterns, interference with daytime functioning, perceived impairment, and distress associated with sleep problems. Each item is rated on a five-point Likert scale (0–4), yielding a total score ranging from 0 to 28. Established interpretive ranges classify scores of 0–7 as no clinically significant insomnia, 8–14 as subthreshold insomnia, 15–21 as moderate clinical insomnia, and 22–28 as severe clinical insomnia^53^. The ISI demonstrates strong content, concurrent, and predictive validity. In this study, it was controlled forthe analyses between insomnia groups.

### Procedure

All participants underwent a clinical interview and screening for confounding sleep disorders (e.g., obstructive sleep apnea, restless legs syndrome, periodic limb movements, REM sleep behavior disorder, narcolepsy). Eligible participants then completed an overnight screening in lab PSG to rule out any primary sleep disorders (other than insomnia). That first PSG night also served as a habituation night.

Participants returned for at least one experimental sleep assessment involving comprehensive PSG. When multiple nights were available (e.g., experimental night and/or intervention vs control night), the control night was selected for analysis. Participants slept in private, sound-attenuated bedrooms, with continuous video and physiological monitoring by a trained research assistant stationed in an adjacent control room.

### Measures

#### Polysomnographic (PSG) Recording

PSG recordings included electroencephalography (EEG), electrooculography (EOG), electromyography (EMG), and electrocardiography (ECG). Respiratory effort (thoracic and abdominal belts), airflow (nasal-oral thermocouple), and oxygen saturation (finger pulse oximetry) were also recorded during the first screening night.

EEG electrodes were positioned according to the international 10–20 system, including at minimum Fz, F3, F4, Cz, C3, C4, Pz, P3, P4, O1, O2, and mastoids (M1–M2). Signals were recorded using SOMNOmedics amplifiers (Somnomedics GmbH, Germany), sampled at 512Hz, referenced online to Pz, and re-referenced offline to the contralateral mastoids (joint M1-M2).

#### Polysomnographic Analyses

Sleep staging (NREM1, NREM2, NREM3, REM, wake) and arousals were scored manually by two independent raters blind to participant group, following standard American Academy of Sleep Medicine^54,55^ criteria, using the Wonambi Python toolbox (https://wonambi-python.github.io)^56^. Sleep macroarchitecture variables were derived from standard scoring procedures and are reported to provide descriptive context for microarchitectural analyses. Extracted sleep parameters included total sleep time (TST), time in bed (TIB), sleep onset latency (SOL), wake after sleep onset (WASO; i.e., how long participants are awake after lights out), sleep stage proportions including % of wake (i.e., how much of the recording is scored as wake), sleep efficiency (SE; i.e., TST/TIB × 100), and sleep fragmentation index (SFI; i.e., # shifts to wake and lighter stages (e.g., NREM1 or NREM2) from NREM3 or REM)/TST [hrs]^57^). Artefacts and poor-quality epochs and electrodes were detected manually.

Oscillatory activities (event-based and spectral power) were automatically analyzed from Fz, Pz, and Cz channels using the seapipe pipeline (https://github.com/nathanecross/seapipe), an open-source Python-based package^58^.

##### Spectral power

EEG spectrum power average (30s of time resolution with artefact excluded) was calculated with a 0.2Hz resolution, by applying a Fast Fourier Transformation (FFT; 50% overlapping, 5s windows, Hanning filter). Mean power was calculated for each 0.25Hz bins between 0.25Hz and 30Hz and for the following frequency bands: SO (0.25-1.25Hz), SWA (i.e., combined delta and SO: 0.25-4Hz), sigma (11.25-16Hz). Given that age differences have been shown to only emerge when isolating SO (<1.25Hz), and as combining SO with delta (1–4Hz) can mask SO-specific alterations^9^, both SO and delta bands were extracted. Both absolute (µV²) and relative power (band/total power; %) were calculated for each band.

##### Event detection

Prior to spindle detection, we performed a data-driven selection of participant-specific sigma-band boundaries using the specparam algorithm to distinguish spectral peaks from aperiodic background activity^59^ within 9-16Hz across combined NREM2 and NREM3 sleep epochs. Such participant-driven sigma peak detection is robust against the inter-individual variability of spindle peak frequencies^60,61^. We observed a typical anteroposterior spindle frequency gradient^62^ with slower sigma peaks on frontal electrodes (Fz) and faster sigma peaks on central and posterior electrodes (Cz, Pz). For each participant, we used the highest peak, in the 9-13Hz range for Fz, 9-16Hz for Cz and 13-16Hz for Pz, to centre the sigma frequency band (with a 4Hz bandwidth) in spindle detection.

Sleep spindles were detected using a consensus-based pipeline designed to reduce algorithm-specific bias by integrating multiple established spindle detection methods within a unified framework^63^.First, we applied 4 validated automatic methods to detect spindles on Fz, Cz, Pz, including algorithms described by Ferrarelli et al, Mölle et al, Ray et al, and Lacourse and colleagues^63–66^. They were used with their described default settings but using the participant-specific sigma frequency band. EEG segments overlapping annotated artefacts or arousals were excluded prior to detection. To obtain a unified set of spindle events, detections from all algorithms were combined using an additive consensus approach. Overlapping detections across algorithms were merged and considered a single spindle event, whereas non-overlapping detections identified by only one algorithm were retained as individual spindles. Consensus spindles failing minimum duration criteria (0.5-3 s) were discarded, and duplicate events were removed. Final consensus spindles were defined as unified spindle events for downstream analyses. The use of an additive consensus threshold was chosen to maximise sensitivity to spindle events while retaining robustness through multi-algorithm detection.

SOs were detected automatically on Fz, Cz and Pz following the procedure proposed by Staresina et al. (2015)^67^. In brief, it involves i) filtering the participant’s SO-band signal (0.5-1.25Hz); ii) identifying the events with a positive-to-negative zero crossing and a subsequent negative-to-positive zero crossing separated by 0.8-2s; iii) retaining the top 25% of events with the largest trough-to-peak amplitudes.

For each participant and electrodes (Fz, Cz and Pz), we extracted SOs and spindle density (number/30s-epoch) across combined NREM2 and NREM3 sleep. Analyses were conducted at Fz as the primary electrode of interest for SWA measures, and both Fz and Cz for spindle measures. Corresponding measures at either Fz or Cz and Pz were included to evaluate the spatial distribution of effects and are described in the Results and Discussion. Fz and Cz findings that are presented in the Tables. Pz findings that are presented in **Supplemental Material**. Relative power spectral is presented. Absolute spectral power as well as spindle and SO amplitude can be found in the **Supplemental Materials**. Tables and figures present estimated marginal means, while raw means for all dependent variables can be found in **Supplemental Table 1, 2, and 3**.

### Statistical Analyses

Sample size estimation was based on Buysse et al. (2008)^68^, who reported a significant group-by-sex interaction in sleep microarchitecture. Accordingly, a minimum of 73 participants (48 with chronic insomnia, 25 healthy sleepers) were targeted. However, G*Power analysis indicated that 180 participants (90 insomnia, 90 healthy sleepers, balanced by sex) would be an adequate sample to detect medium-sized effects (f = .25) with 80% power at α = .05. Statistical analyses were conducted using SPSS 31.0.0.0. Participants were considered outliers if they showed extreme values (±3.29 SD) on three or more dependent variables; this conservative approach ensured cases were not excluded due to chance variation on a single measure.

Assumptions of normality and homogeneity of variance were assessed using Shapiro–Wilk tests, skewness and kurtosis z-scores (±1.96), and Levene’s tests. Several dependent variables showed deviations from these assumptions; therefore, we used Generalized Linear Models (GLMs) with Huber–White/Sandwich robust estimators of variance and bootstrapped (1,000 samples) bias-corrected 95% confidence intervals to provide robust inference under unequal variances. GLM models included two categorical factors (Sex: female/male; Group: insomnia/healthy sleepers). Group × Sex interactions were examined, along with main effects of Sex and Group controlling for age. Age was entered as a continuous covariate to adjust for potential differences in age distribution between groups. This approach enabled us to test age effects as well as age-adjusted main effects of Sex, and Group on brain oscillations. All analyses used an α level of 0.05. Bootstrapped confidence intervals and Benjamini–Hochberg FDR correction were applied across analyses to account for multiple comparisons^69^. Analyses used listwise deletion for missing data. Together, these methods provide robust parametric inference while mitigating the influence of non-normality and heteroscedasticity.

## Results

### Demographics

The final sample (N = 222, age range 18-82) included 119 individuals with chronic insomnia (INS; 54%) and 103 healthy sleepers (HS; 46%). The sample comprised 67% females (*n* = 148), with 33% males (*n* = 74). When stratified by both sleep group and biological sex, the sample comprised 34 male INS (MINS; 15%), 85 female INS (FINS; 38%), 40 male HS (MHS; 18%), and 63 female HS (FHS; 29%). This distribution of higher representation of females in both the INS and HS groups is consistent with prior findings of sex distributions in insomnia research^26,41^.

There was a statistically significant age difference across all four groups (Wald χ²(3) = 38.79, p < .001; **Supplemental Table 4**). Bootstrapped pairwise comparisons indicated that FINS were older than FHS (β = 13.99, p < .001), MINS (β = -8.47, p = .005), and MHS (β = 15.77, p < .001). MHS were also significantly younger than MINS (β = 7.29, p= 0.04). Female and male HS did not differ in terms of age. Hence, all analyses were controlled for age.

Within the INS group, insomnia severity (based on the ISI^53^) ranged from 8 (mild) to 27 (severe) with an average moderate severity score (16.56 ± 3.96). There was a significant difference between FINS and MINS (Wald χ²(1) = 5.65, p= .017), with females reporting more severe insomnia (17.09 ± 0.42) than males (15.24 ± 0.66, β = -1.86, p < .05). Insomnia severity did not differ by age (Wald χ²(1) = 0.01, p= .942).

### Sleep Macroarchitecture

Descriptive sleep macroarchitecture for the sample is presented in **Supplemental Tables 1-4**. These metrics provide context for subsequent analyses of NREM oscillatory activity. No hypotheses were tested for these variables.

#### Age Effect in Sleep Macroachitecture Across All Participants

Increased age was associated with a significant reduction of TST, SE, and time spent in NREM2, NREM3, and REM. Increases in time awake, WASO, and sleep fragmentation were also observed with increased age, but not TIB, SOL, or time spent in NREM1 (all p > 0.05; **Supplemental Table 1**).

#### Effect of Sleep Group on Sleep Macroachitecture

After controlling for age, INS spent significantly more time awake, WASO, and less time in NREM1, NREM2, NREM3, REM, TST, and SE compared to HS. However, TIB, SOL, and sleep fragmentation were not impacted by the presence of insomnia (all p > 0.05; **Supplemental Table 2**).

#### Effect of Biological Sex on Sleep Macroachitecture

While age was controlled, males had more time spent in NREM1, had more WASO, and sleep fragmentation compared to females. Sex did not significantly impact other sleep architecture variables (i.e., time spent awake, in NREM2, NREM3, REM, TIB, TST, SOL, and SE; all p > 0.05; **Supplemental Table 3**).

#### Group by Sex Interaction on Sleep Macroachitecture

After controlling for age, we found group by sex differences in measures of sleep macro-architecture (**Supplemental Table 4**). Both HS groups had higher TST, SE, time in NREM1, NREM2, NREM3 and REM. They also had lower WASO, and time awake compared with both INS groups. There were no group differences in SOL. Sleep fragmentation differed significantly between all 4 groups and appeared to mainly be driven by sex (Wald χ²(3) = 27.13, p < .001) with males’ sleep appearing more fragmented than females. FINS showed a reduced NREM1, NREM2, NREM3, and REM percentage relative to FHS. The only significant sex difference within insomnia groups was for percentage of time spent in NREM3, where MINS displayed less NREM3 compared to FINS (β = 6.74, 95% CI [3.58, 9.92]).

### NREM Sleep Microarchitecture

#### Age Effect in Measures of Spindle and SWA Across All Participants

In Cz, increasing age predicted significant reductions in spindle density (β = –0.02, p < .001, 95% CI [–0.025, –0.013]) and in Fz for relative SWA (β = –0.001, p < .001, 95% CI [–0.002, –0.001]). SO density and relative sigma power were preserved across age in all channels (**Table 1**). The main effects of age were seen across electrodes (**Supplemental Table 8**).

**Table 1.** - Age Differences in Measures of Spindle and SWA Across All Participants.

| Fz) | Wald $\chi^2$ (df= 1) | p | Coefficient | SE | p | 95% Lower | Upper | Difference |
| --- | --- | --- | --- | --- | --- | --- | --- | --- |
| Event Detection |  |  |  |  |  |  |  |  |
| Spindle Density | 41.55 | < .001 | **-0.02** | **0.003** | **<.001** | **-0.022** | **-0.012** | ↓ |
| SO Density | 0.63 | .427 | -0.009 | 0.011 | .444 | -0.032 | 0.012 | - |
| Power Spectral Analysis |  |  |  |  |  |  |  |  |
| Relative Sigma power | 2.67 | .102 | $5.70 \times 10^{-5}$ | $3.59 \times 10^{-5}$ | .113 | $-6.10 \times 10^{-6}$ | 0.000 | - |
| Relative SWA | 33.24 | < .001 | **-0.001** | **0.000** | **<.001** | **-0.002** | **-0.001** | ↓ |
| Cz) |  |  |  |  |  |  |  |  |
| Event Detection |  |  |  |  |  |  |  |  |
| Spindle Density | 43.78 | < .001 | **-0.02** | **0.003** | **<.001** | **-.025** | **-.013** | ↓ |
| SO Density | 1.47 | .225 | -0.014 | 0.011 | .222 | -0.037 | 0.008 | - |
| Power Spectral Analysis |  |  |  |  |  |  |  |  |
| Relative Sigma power | 7.21 | .007 | 0.000 | $1.09 \times 10^{-6}$ | .008 | $3.58 \times 10^{-5}$ | .000 | - |
| Relative SWA | 36.75 | < .001 | **-0.002** | **0.000** | **< .001** | **-.002** | **-.001** | ↓ |

Note. Bias-corrected confidence intervals are presented. *p* = adjusted p. Significant results are bolded. Sigma power (11.25-16Hz), SWA (0.25-4Hz), SO power (0.25-1.25 Hz), NREM2 + NREM3

#### Effect of Sleep Group on Measures of Spindle and SWA

After controlling for age, INS showed significantly lower central spindle density (β = –0.80, p < .001, 95% CI [–1.00, –0.60]), and frontal SO density (β = –2.72, p <.001, 95% CI [–3.52, –2.01]) compared to HS (**Table 2**). This effect was seen across all electrodes. Groups did not differ in central sigma power or frontal relative SWA across electrodes (**Supplemental Table 9**). However, frontal relative sigma power (β = -0.01, p < .001, 95% CI [-0.01, -0.007]) was lower in INS than HS.

**Table 2.** - Age Controlled Group Differences in Measures of Spindle and SWA.

| Fz) | INS (n= 119) | HS (n= 103) | Wald $\chi^2$ (df= 1) | p | Coefficient | SE | p | 95% Lower | Upper | Difference |
| --- | --- | --- | --- | --- | --- | --- | --- | --- | --- | --- |
| Event Detection |  |  |  |  |  |  |  |  |  |  |
| Spindle Density | 2.13 ± .05 | 2.91 ± .08 | 74.27 | < .001 | **-0.77** | **0.09** | **< .001** | **-0.96** | **-0.60** | **INS < HS** |
| SO Density | 5.12 ± .14 | 7.85 ± .37 | 49.61 | <.001 | **-2.72** | **0.399** | **<.001** | **-3.52** | **-2.01** | **INS < HS** |
| Power Spectral Analysis |  |  |  |  |  |  |  |  |  |  |
| Relative Sigma Power | .02 ± .001 | .03 ± .001 | 50.63 | <.001 | **-0.01** | **0.001** | **<.001** | **-0.01** | **-0.007** | **INS < HS** |
| Relative SWA | .79 ± .01 | .79 ± .01 | .125 | .723 | -0.003 | 0.008 | .731 | -0.018 | 0.013 | - |
| Cz) |  |  |  |  |  |  |  |  |  |  |
| Event Detection |  |  |  |  |  |  |  |  |  |  |
| Spindle Density | 2.33 ± .06 | 3.13 ± .09 | 59.10 | <.001 | **-0.80** | **0.102** | **<.001** | **-1.00** | **-0.60** | **INS < HS** |
| SO Density | 5.04 ± .14 | 7.72 ± .38 | 47.51 | <.001 | **-2.68** | **0.390** | **<.001** | **-3.45** | **-1.92** | **INS < HS** |
| Power Spectral Analysis |  |  |  |  |  |  |  |  |  |  |
| Relative Sigma Power | .03 ± .001 | .027 ± .001 | 5.17 | .023 | 0.004 | 0.002 | .027 | 0.00 | 0.008 | - |
| Relative SWA | .75 ± .01 | .76 ± .01 | 0.51 | .477 | -0.006 | 0.008 | .483 | -0.022 | 0.010 | - |

Note. Values are reported as estimated marginal means (age controlled) ± standard error with bias-corrected confidence intervals. *p* = adjusted p. Significant results are bolded. Sigma power (11.25-16Hz), SWA (0.25-4Hz), SO power (0.25-1.25 Hz), NREM2 + NREM3, INS = insomnia group; HS = healthy sleeper group.

#### Effect of Biological Sex on Measures of Spindle and SWA

After controlling for age, frontal relative SWA (β = –0.02, p = .032, 95% CI [–0.038, -0.003]) was lower in males than females, as well as posterior (**Table 3 and Supplemental Table 10**). No significant sex differences were found for central spindle density (β = –0.10, p = .329, 95% CI [–0.29, 0.10]) or frontal SO density (β = –0.09, p = .836, 95% CI [–0.951, 0.703]) across electrodes (**Supplemental Table 10**). Central relative sigma power was equal between sexes (β = 0.003, p = .135, 95% CI [-0.001, 0.007]), but frontal relative sigma power (β = 0.004, p = .007, 95% CI [0.002, 0.007]) was higher in males than in females. Similar results were found in Pz (**Supplemental Table 10**).

**Table 3.** - Age Controlled Sex Differences in Measures of Spindle and SWA.

| Fz) | Male (n= 74) | Female (n= 148) | Wald $\chi^2$ (df= 1) | p | Coefficient | SE | *p* | 95% Lower | Upper | Difference |
| --- | --- | --- | --- | --- | --- | --- | --- | --- | --- | --- |
| Event Detection |  |  |  |  |  |  |  |  |  |  |
| Spindle Density | 2.44 ± .07 | 2.60 ± .05 | 3.19 | .074 | -0.16 | .09 | 0.065 | -0.34 | 0.004 | - |
| SO Density | 6.44 ± .34 | 6.53 ± .23 | .048 | .827 | −0.09 | .409 | .836 | −0.951 | 0.703 | - |
| Power Spectral Analysis |  |  |  |  |  |  |  |  |  |  |
| Relative Sigma power | .03 ± .001 | .02 ± .001 | 9.30 | .002 | **0.004** | **.001** | **.007** | **0.002** | **0.007** | **M > F** |
| Relative SWA | .78 ± .01 | .80 ± .004 | 5.49 | .019 | **−0.02** | **.009** | **.032** | **−0.038** | **-0.003** | **M < F** |
| Cz) |  |  |  |  |  |  |  |  |  |  |
| Event Detection |  |  |  |  |  |  |  |  |  |  |
| Spindle Density | 2.68 ± .08 | 2.78 ± .06 | 0.94 | .333 | −0.10 | .101 | .329 | −0.29 | 0.10 | - |
| SO Density | 6.31 ± .34 | 6.45 ± .24 | 0.12 | .733 | −0.14 | .423 | .738 | −0.98 | 0.70 | - |
| Power Spectral Analysis |  |  |  |  |  |  |  |  |  |  |
| Relative Sigma power | .03 ± .002 | .03 ± .001 | 2.24 | .134 | 0.003 | .002 | .135 | −0.001 | 0.007 | - |
| Relative SWA | .75 ± .01 | .76 ± .004 | 4.35 | .037 | −0.018 | .009 | .048 | −0.036 | 0.000 | - |

Note. Values are reported as estimated marginals means (age controlled) ± standard error with bias-corrected confidence intervals. *p* = adjusted p. Significant results are bolded. Sigma power (11.25-16Hz), SWA (0.25-4Hz), SO power (0.25-1.25 Hz), NREM2 + NREM3, M = male; F = female.

#### Group by Sex Interaction on Measures of Spindle and SWA

Concerning sigma power, we found a Group-by-sex interaction in central sigma power (Wald χ²(3) = 9.68, p = .022; **Table 4**). FINS showed lower central relative sigma power than MINS (β = -0.006, 95% CI [-0.001, -0.012], p = .030). This relationship was maintained even after controlling for ISI score (β = 0.005, p = .014) and was also found in Fz, but not Pz (**Supplemental Table 11, 12, 13**). MINS displayed higher central relative sigma power compared to MHS group (**Table 4 and Figure 1B**). Within the HS group, FHS exhibited lower posterior relative sigma power than MHS (β = 0.006, 95% CI [0.001, 0.011], p = .025), but not in Fz or Cz (all p > 0.05; **Supplemental Table 11, 12, 13**).

**Figure 1.**
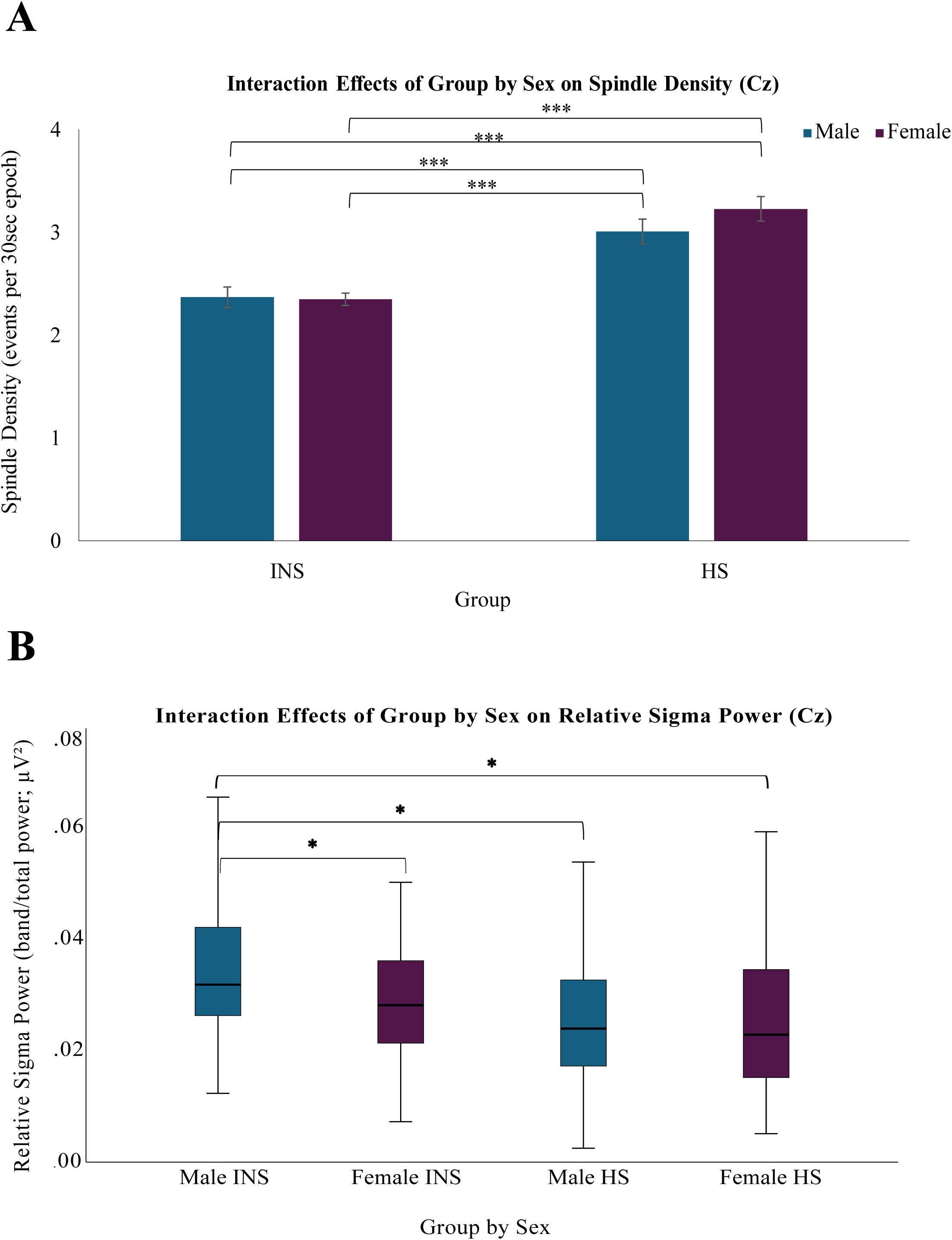
Significant Group by Biological Sex Interaction on Spindle Activity. A) *Estimated marginal means ± SE from the generalized linear model of spindle density (detected on Cz) per Group and Sex* B) *Estimated marginal means ± SE from the generalized linear model of relative sigma power (detected on Cz; 11.25-16Hz) per Group and Sex* * p < 0.05 ** p < 0.01 *** p < 0.001 + p > .05 but became significant after bias-corrected confidence intervals

**Table 4.**
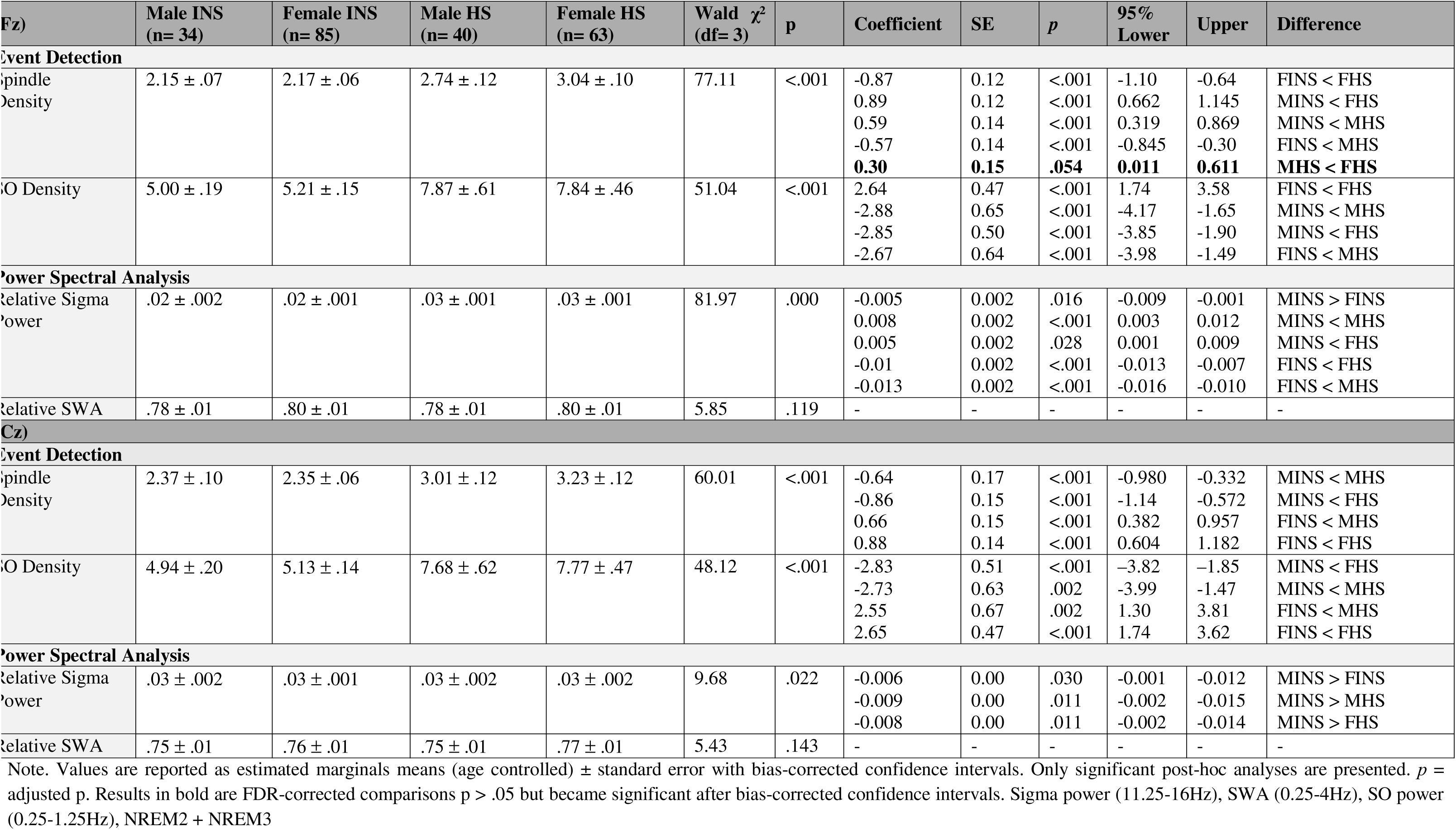
- Age Controlled Group by Sex Differences in Spindle and SWA Measures.

For spindle density, we found a Group-by-sex interaction (Wald χ²(3) = 77.11, p < .001) that was driven by MHS exhibiting less frontal spindle density than FHS (β = 0.30, 95% CI [0.011, 0.611], p = .054), but not central or posterior spindle density (**Table 4**). While there were no sex differences within the INS group, both FINS and MINS exhibited lower central spindle density relative to the HS sex counterparts (FINS: β = 0.88, p < .001; MINS: β = -0.64, p < .001; **Table 4 and Figure 1A**). The same relationship was seen for frontal and parietal spindle density **Supplemental Table 11, 12, 13)**.

For SWA, after adjusting for age, there were no Group-by-sex interactions in relative SWA in Fz (*Wald χ²*(3) = 5.85, *p* =.119) or across other electrodes (**Table 4 and Supplemental Table 11, 12, 13)**. There was, however, Group-by-sex differences in SO density within frontal (*Wald χ²*(3) = 51.04, *p* < .001), central and parietal. There were no differences between either sex with INS on SO density across all electrodes. Relative to the HS sex counterparts, both FINS and MINS exhibited lower frontal SO density (FINS: β = 2.64, p < .001; MINS: β = -2.88, p < .001; **Table 4 and Figure 2**).

**Figure 2.**
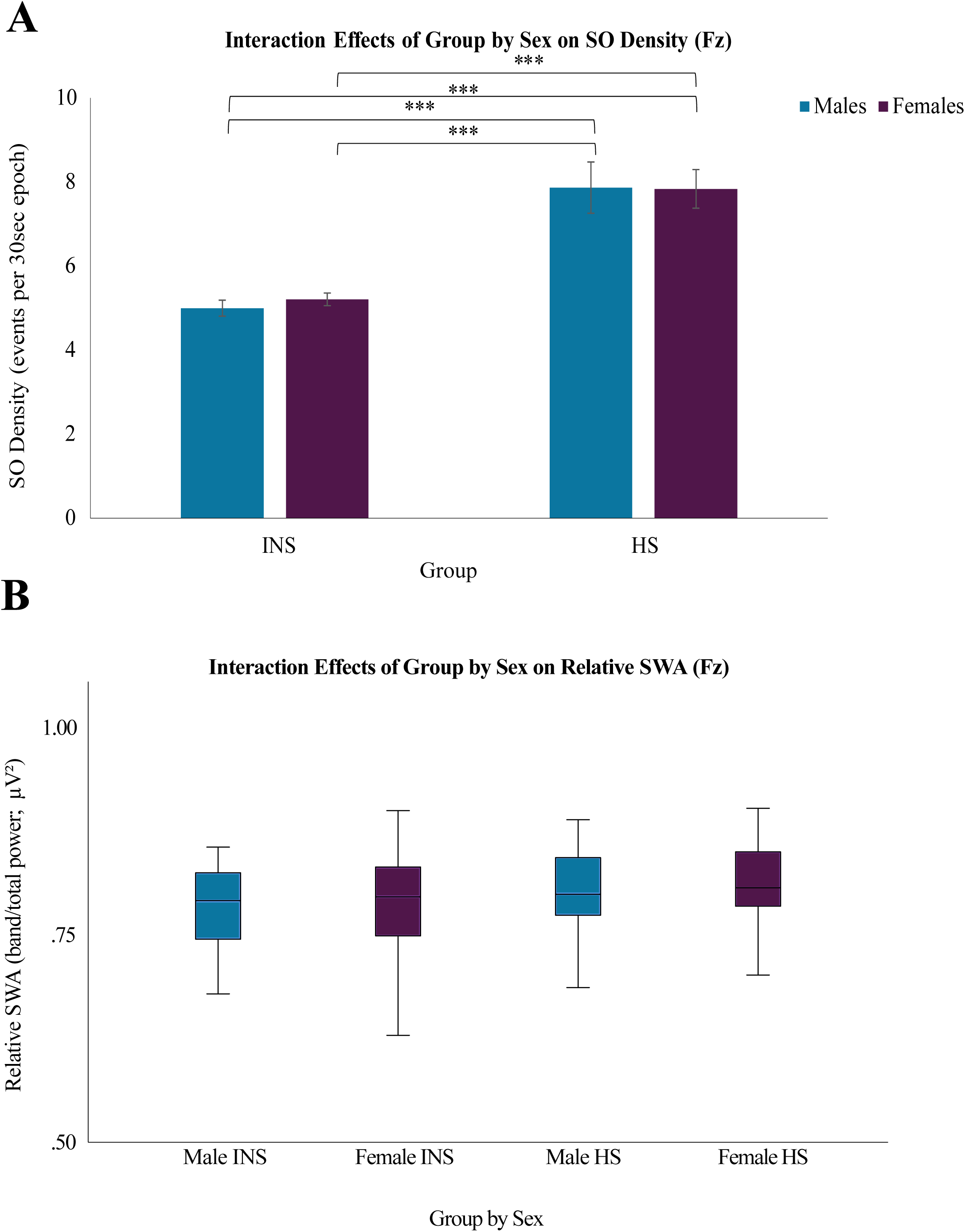
Significant Group by Biological Sex Interaction on SWA. A) *Estimated marginal means ± SE from the generalized linear model of SO density (detected on Fz; 0.25-1.25 Hz) per Group and Sex* B) *Estimated marginal means ± SE from the generalized linear model of relative SWA (detected on Fz; 0.25-4 Hz) per Group and Sex* * p < 0.05 ** p < 0.01 *** p < 0.001

See a summary of the results in **Table 5**. Additional group by sex analyses on spindle and SO amplitude, SO power, and absolute sigma and SWA can be found in **Supplemental Material.**

**Table 5.** - Summary of Results.

| Outcome | Age Effects | Sex Effects* | Insomnia Effects | Key Interactions | Primary Drivers |
| --- | --- | --- | --- | --- | --- |
| **Insomnia Severity** | X | Females > Males | — | Female INS > Male INS | **Sex** |
| **Sleep Architecture** |  |  |  |  |  |
| NREM1 | X | Females < Males | X | Female INS < Female HS<br>Both INS groups < than sex-matched HS | **Sex, Sex × Insomnia** |
| NREM2, NREM3, REM | Declined with age | X | INS < HS | Female INS < Female HS<br>Both INS groups < than sex-matched HS<br>Female INS > Male INS (in NREM3 only) | **Age, Insomnia, Sex × Insomnia** |
|  | Declined with age | X | INS < HS | Both INS groups < than sex-matched HS | **Age, Insomnia** |
|  | Increased with age | Females < Males (just WASO) | INS > HS | Both INS groups > than sex-matched HS | **Age, Insomnia** |
|  | Increased with age | Females < Males | X | SFI driven by sex | **Age, Sex** |
|  | X | X | X | X | X |
| **Relative Sigma Power Fz** | X | Females < Males | INS < HS | Females INS < Male INS<br>Both INS groups < than sex-matched HS | **Sex, Insomnia, Sex × Insomnia** |
| **Relative Sigma Power** | X | X | X | Females INS < Male INS<br>Male INS > Male HS | **Sex × Insomnia** |
| **Spindle Density Fz** | Declined with age | X | INS < HS | Both INS groups < than sex-matched HS | **Age, Insomnia** |
| **Spindle Density Cz** | Declined with age | X | INS < HS | Both INS groups < than sex-matched HS | **Age, Insomnia** |
|  | Declined with age | Females > Males | X | X | **Age, Sex** |
|  | X | X | INS < HS | Both INS groups < than sex-matched HS | **Insomnia** |

\*Combined INS and HS groups, — Not examined, X no effect, INS= insomnia group, HS= healthy sleeper group, TIB = time in bed; TST = total sleep time; SOL = sleep onset latency; WASO = wake after sleep onset; SE= Sleep efficiency; NREM = non-rapid eye movement sleep; REM = rapid eye movement sleep; SFI = sleep fragmentation index;

## Discussion

### Summary

This study investigated how age, the presence of chronic insomnia disorder and biological sex, individually and jointly, influence key NREM oscillatory markers of sleep quality. We tested whether sex and insomnia exert effects beyond aging, and whether females with insomnia, who typically report greater symptom severity, show disproportionate alterations in NREM microarchitecture. Age, insomnia, and sex each contributed uniquely to SWA and spindle activity. Insomnia primarily drove reductions in spindle and SO density above and beyond age or sex effects, while sex accounted for differences in relative SWA. While relative SWA was higher in females across the full sample, sex differences were not significant within insomnia. Both insomnia and sex influenced central relative sigma power, with females showing lower sigma than males experiencing insomnia. In addition, both females and males with insomnia exhibited deficits in most measures compared to healthy sleepers. These patterns underscore that, independent of age, both sex and insomnia uniquely shape NREM oscillatory activity, and are dependent on the specific measures used.

### The Impact of Age on NREM Neural Activity

In our sample, age selectively affected certain brain oscillations, while other NREM features were resilient. Specifically, typical age-related reductions were present in Cz for spindle density, and in frontal, central and parietal regions for relative SWA, likely due to structural and functional alterations in thalamocortical networks^7,23,31,70,71^, as well as cortical thinning and diminished large-scale neural synchrony as a natural process of aging^72–74^. The frontal SWA decline is consistent with known prefrontal cortical thinning, whereas the parietal SWA reduction may reflect broader network-level changes in NREM synchrony. Interestingly, the Age effect on spindle density was restricted to Cz, which may relate to our participant-specific sigma-band detection approach. The adapted frequency range at Cz likely captured both slow and fast spindles, thereby reflecting a more general spindle activity measure. In contrast, the adapted detection ranges at frontal and parietal electrodes may have preferentially targeted slow (Fz) and fast (Pz) spindles, respectively. Age-related changes in general spindle measures at central sites may therefore not extend to specific fast or slow spindle effects. However, relative sigma power as well as SO density in Cz appeared fairly consistent across adulthood, and were more selectively influenced by insomnia and/or sex, contrasting reports of strong age-related declines in spindle and general measures of SWA^31,73,75^. Such discrepancies may reflect differences in sample composition and methodology such as the oscillatory feature and its detection method (e.g., spectral power or event detection). Many previous studies did not explicitly control for disturbed sleep or sex differences, factors that our results show can influence spindle and SWA measures^73^. Additionally, our use of participant-specific sigma-band detection, combined with averaging across NREM2 and NREM3, likely mitigated some age-related declines that may have been overestimated in prior studies using fixed frequency bands or absolute power measures. This approach, together with separate analyses of frontal and centro-parietal spindles, may explain why we observed selective age effects rather than widespread age-related reductions.

### The Impact of Insomnia on NREM Neural Activity

Within Fz, relative SWA appeared largely preserved in insomnia, whereas SO and spindle density, and relative frontal sigma power (but not central) emerged as key markers of sleep quality that were disrupted by insomnia disorder. This pattern highlights distinct measure-specific effects, emphasizing how event-based and spectral indices capture complementary aspects of cortical synchronization and NREM homeostasis. Insomnia was the primary driver of reductions in SO density, consistent with hyperarousal and shallower sleep depth^6,76^ in insomnia, a pattern further supported by elevated sleep fragmentation indices and reduced NREM3 in our insomnia sample. In contrast, relative SWA was not impacted by insomnia, aligning with prior work indicating that insomnia-related SWA deficits may be more reliably detected in SO-specific measures^8,9^ rather than broadband delta power^8,9,26,68,77–79^. In fact, delta power can decrease^24^, remain relatively stable or even increase during sleep deprivation even when other markers of sleep depth are disrupted^80^. Event-based measures of low-frequency (0.25-1.5 Hz) cortical synchrony, such as SO density, may be particularly sensitive to sleep fragmentation and hyperarousal^81^ that are often present in individuals with insomnia. This is because coordinated activity across widespread cortical networks are strongly influenced by arousal-related subcortical structures such as the thalamus^82^. In contrast, SWA is a broader and less temporally specific measure, averaging spectral power (up to 40Hz) across entire epochs. As a result, SWA can appear relatively preserved even when the fine-grained frequency or organization of individual SO events is disrupted^80^. It is also quite possible that certain insomnia subtypes (e.g., insomnia with physiological hyperarousal) may disrupt slow-wave microstructure, without altering global measures of homeostatic sleep regulation^8,23,68,77,83^.

SOs also represent coordinated switches between cortical states that influence the timing and expression of other sleep rhythms, and play a critical role in coordinating spindles, supporting memory consolidation, and facilitating cellular recovery^81^. Therefore, it is unsurprising that the presence of insomnia was also associated with robust and consistent alterations in spindle characteristics. These findings align with prior research^26,84,85^ and studies demonstrating that reduced spindle activity is a reliable marker of disrupted sleep^12,22,23^. In this context, insomnia emerged as the primary driver of reductions in frontal and central spindle density and contributed to decreases in relative frontal sigma power, underscoring the particular vulnerability of spindle-generating mechanisms in insomnia. Notably, studies reporting reductions in spindle activity similar to those observed here employed frontal and central EEG electrodes and were of good methodological quality^84,85^. Although findings regarding spindle impairment in insomnia have been mixed, this variability is likely attributable to methodological heterogeneity^86^ and the complexity of the disorder. Specifically, differences in study design may explain some inconsistencies: for example, some studies have found increased sigma power in insomnia^77,87^ but these findings were often limited by factors such as narrow age ranges, restricted spindle frequency bands (e.g., 12.5–16 Hz) and localization exclusively to the left central region (e.g., C3), which reduces sensitivity to frontal spindles. Despite these differences, our results are consistent with broader literature highlighting the role of spindles in promoting sleep stability, and buffering stress^11,88,89^. Moreover, they support evidence that spindle activity may serve as a useful predictor of response to Cognitive Behavioural Therapy for Insomnia^90^(CBTi; the gold standard treatment for insomnia^91^).

### The Impact of Sex on NREM Neural Activity in Insomnia

Contrary to much of the existing literature in healthy populations^26,29,92^, we did not observe sex differences in spindle density or SO density in any group. This was unexpected, as females have been shown to exhibit higher spindle density and, in some studies, greater SWA^26^. One possibility is that sex differences in event-based measures are partially contingent on age distribution, hormonal status, or topographic specificity, factors that may differ across studies. In this context, the absence of spindle and SO density differences in our sample suggests that sex effects in discrete oscillatory events may not be as robust across samples as spectral findings.

Indeed, we found that relative frontal sigma power and SWA emerged as the primary sleep features differentiating the sexes, with sex exerting a primary influence on relative SWA. Although healthy males and females as well as male sand females with insomnia did not differ in relative SWA, a sex differences emerged only when sleep groups were combined. Males showed lower relative SWA overall, consistent with findings on absolute SWA^29,48,93–102^, but not relative delta power^95,97,98,103,104^. When examined within groups, reduced sample size may have limited statistical power to detect small effects. Collapsing across sleep groups increases power and may have allowed a subtle overall sex difference in SWA to become detectable. Importantly, this pattern suggests that sex differences in SWA are not uniquely driven by healthy sleep or preserved in insomnia per se but instead reflect a modest global shift that becomes visible only when statistical power permits.

#### Hormonal Modulation of Sigma Activity: Implications for Female Sleep Stability

In insomnia, the typical female advantage in sigma power^26^ observed in healthy sleepers disappeared with females exhibiting lower relative frontal and central sigma power than males, supporting the interpretation that insomnia may selectively disrupt mechanisms supporting female sleep stability. Although age was statistically controlled for, the average age of the female insomnia sample fell within the peri- to postmenopausal range (>45 years old), when there is progressive ovarian follicle loss and reduced ovarian responsiveness^105^. Estradiol and progesterone levels decline, and residual hormonal variability during this transition has been linked to difficulties initiating sleep and increased nocturnal awakenings^106–108^. These hormonal changes may be particularly relevant for spindle activity, given the neurophysiological mechanisms underlying their generation. Sleep spindles arise from reciprocal interactions between inhibitory neurons (e.g., GABA) in the thalamic reticular nucleus interacting with excitatory thalamocortical neurons (e.g., glutamate), producing the rhythmic activity that travels through thalamocortical circuits^34^. Estradiol and progesterone are thought to modulate GABAergic and serotonergic signaling, thereby supporting thalamocortical excitability and spindle generation^109^. Consistent with hormonal fluctuation, spindle density has been shown to increase during the luteal phase and with oral contraceptive use during the reproductive years as well as menopausal hormone therapy^110,111^ likely via progesterone-enhanced GABAergic transmission within cortical and thalamic networks. Given the central role of spindles in maintaining sleep continuity and depth^19–21^, reductions in sigma activity may represent a key mechanism contributing to insomnia vulnerability^9–11^, and overall higher prevalence and severity rates of insomnia in females. Accordingly, the lower sigma power observed in females with insomnia may reflect the combined influence of female sex-steroid hormones on thalamocortical networks and the clinical expression of sleep disruption, highlighting a plausible mechanistic pathway underlying sex-specific vulnerability to insomnia.

In contrast to the sex-specific effects observed for relative sigma power, no differences between males and females with insomnia were observed for frontal or central spindle density, SO density, or relative SWA. To our knowledge, no previous studies have explicitly examined sex differences or sex-by-group interactions in spindle or SO density in insomnia^26^. For spindle density in Fz and Cz, both insomnia groups showed lower values than healthy sleepers, suggesting that insomnia broadly affects spindle generation but that the sex-specific vulnerability in females may be most apparent in sigma power rather than event counts. Prior literature supports that spectral sigma power and spindle density do not always correlate^62,86^. Sigma power reflects the overall energy in the sigma frequency range, whereas spindle density quantifies only the number of clearly defined spindles. Functional studies of developmental sleep^112^, and other sleep disorders^86,113^ have similar divergences. Specifically, sigma power correlates with aging and cognitive performance, but spindle density does not systematically, indicating that these measures capture overlapping but distinct aspects of thalamocortical activity.

The absence of sex differences in relative SWA among individuals with insomnia also aligns with the limited existing literature using both relative and absolute measures^68,77,79,87,114,115^. From a mechanistic perspective, this pattern indicates that while thalamocortical sigma activity may be sensitive to sex-specific processes, such as hormonal fluctuations, the cortical networks generating SO and delta activity seem to be similarly vulnerable to the effects of insomnia across sex. This may be alternatively attributable to a potential compensatory neuroadaptive mechanisms that help maintain sleep homeostasis in the face of gradual hormonal changes.

These findings demonstrate that spindle and SWA metrics do not always change in parallel, highlighting the value of integrating both event-based and spectral measures, as each provides distinct and complementary insights into how insomnia and sex shape NREM sleep physiology that a single metric alone might miss.

#### Insomnia-Related Sleep Changes in Females

Some deficits in spindle and SWA measures were apparent only when females with insomnia were compared to female healthy sleepers, indicating that insomnia may involve deviations from normative female sleep patterns, such as reduced spindle and SO density. This finding is consistent with the higher insomnia severity reported by females in our sample, suggesting that their subjective complaints may reflect disruptions relative to typical female sleep, which can be missed if studies simply compare females to male insomnia profiles. Nevertheless, longitudinal research is required to determine the emergence of these alterations, whether they develop as a consequence of chronic insomnia, or contribute to subjective ratings of sleep quality, and the extent to which they persist over time.

### Methodological Impacts on Sex Differences in Insomnia

Our results revealed both region-specific and global robust patterns across the scalp in how sex and insomnia affect NREM sleep. Sleep group and sex effects in spindle, SO density, and relative SWA were consistent across electrodes. These findings indicate that insomnia-related and sex-related changes in oscillatory activity, were widespread across the cortex rather than restricted to particular regions, suggesting these effects reflect global alterations in oscillatory activities. Frontal channels were particularly sensitive to sigma activity in insomnia, while frontal and posterior channels were sensitive to sex differences in sigma activity and relative SWA. Lastly, frontal and central channels were particularly sensitive to sigma power, with male insomnia participants showing higher frontal and central sigma than female insomnia participants, reflecting more global thalamocortical network activity impacts. Overall, these findings highlight that while some sleep features are regionally specific, others are robust across the scalp, emphasizing the importance of both electrode coverage and individualized analysis in insomnia research.

Finally, our findings highlight important distinctions between absolute and relative EEG power when interpreting sex differences in insomnia. Consistent with systematic reviews^24^, which suggested that relative power measures may be more sensitive at detecting power spectral differences during NREM sleep in those with insomnia, we observed that sex differences within insomnia varied depending on the power metric used. For SWA, females with insomnia exhibited higher absolute power than males, but not relative power, suggesting that absolute measures may be further confounded by sexually dimorphic anatomical factors such as skull thickness and tissue conductivity^73,94,116,117^, rather than true differences in frequency-specific neural processes. In contrast, sigma power showed more nuanced patterns: relative frontal sigma power was lower in females with insomnia compared with males but showed the opposite relationship for posterior absolute power. Taken together, these results suggest that absolute power reflects global amplitude differences influenced by anatomy and age, while relative power, which normalizes power within each individual, may provide a more sensitive and physiologically meaningful index of sex-specific alterations in insomnia.

### Strengths and Limitations

Our study has several methodological strengths that enhance the reliability and validity of the observed patterns. First, we conducted high-quality and low risk of bias research^118^, evident by the large, justified sample size, rigorous ascertainment of exposure (i.e., diagnosis via structured clinical interview, PSG screening for comorbid sleep disorders, and use of validated clinical measures), careful comparability of groups, objective and validated outcome assessment, and appropriate statistical testing. We also used a merged spindle-detection approach from validated detectors^63–66^, reducing the bias inherent in any single algorithm and maximizing the robustness of event identification. Detection of brain oscillations was further optimized by applying adapted frequency bands, allowing the algorithm to accommodate well-established inter-individual variability^119^. Importantly, we examined relative and absolute power to understand true differences in neural activity^116,117^. Lastly, this study was the first to examine sex differences in individuals with insomnia across spindle and SO events, and their sleep group × sex interactions.

Several limitations of our study warrant acknowledgement. First, even though statistical models adjusted for unequal group sizes, the number of males with insomnia and healthy sleepers was smaller than females, which may limit precision for sex-specific estimates. However, the present sample constitutes one of the largest available cohorts including males with insomnia and healthy sleepers^26^, strengthening the interpretability. Second, the sample was drawn from multiple studies that employed harmonized methodology but differed in age-related inclusion criteria, introducing potential heterogeneity in lifespan-related effects. This heterogeneity also represents a strength, allowing examination of oscillatory activity across a broad adult age range, and age was explicitly modeled to reduce confounding. Additionally, one of the four contributing insomnia datasets included participants with comorbid anxiety or depression. While this limits causal attribution, psychiatric comorbidity is common in chronic insomnia and thus enhances ecological validity. Moreover, medication use was carefully controlled across studies, reducing the likelihood that observed effects were driven by pharmacological influences. Lastly, this study did not have information on menstrual cycle phase, menopausal status or hormonal dosage, all of which may have improved our understanding of sex differences in brain oscillations. Future longitudinal and multimodal studies integrating additional oscillatory and clinical measures will be critical for clarifying the mechanisms responsible for these complaints

### Clinical and Translational Relevance

These findings have important clinical and translational implications for improving the prevention and treatment of chronic insomnia. As insomnia is more prevalent in females and contributes substantially to lost productivity, worsening mental and physical health, and health-care utilization^120,121^, identifying the mechanisms behind sex- and age-specific vulnerabilities is valuable. By demonstrating that NREM oscillatory features vary systematically by age, sex, and insomnia status, this study highlights physiological markers that could help personalize care. For instance, lower spindle density has been found to predict poorer response to CBT I ^90^. Therefore, individuals with insomnia who show impaired spindle synchrony (patterns that appear more common in females based on our current findings), may have an increase vulnerability that maintains symptoms^89^, and may not experience sufficient physiological change from treatments that primarily target behavioral and cognitive contributors to hyperarousal^122^ rather than the underlying oscillatory mechanisms. This subgroup may require CBT-I augmented with physiology-focused interventions that downregulate arousal, strengthen sleep stability, and meaningfully influence neural activity (e.g., exercise, TMS, rocking stimulation)^83,123–129^. These profiles therefore offer a pathway toward biomarker-informed treatment matching rather than relying solely on subjective symptom reports. Understanding these sex- and age-dependent mechanisms also supports the adaptation of preventive strategies that address biopsychosocial contributors to insomnia (such as genetic risk, caregiving stress, hormonal transitions, and higher rates of anxiety and depression in females)^130–132^ ultimately strengthening efforts to promote sleep health across the lifespan^133^.

### Conclusion

Taken together, our findings indicate that, beyond well-established age effects, meaningful sex differences exist in insomnia-related NREM oscillatory activity. These results highlight the need for sleep research and clinical practice to adopt sex- and gender-informed methodologies and carefully stratify samples by age. Future studies should explicitly examine sex-specific and insomnia-related effects on sleep microarchitecture, including meta-analytic approaches to strengthen the evidence base. An important next step is to determine whether first-line treatments such as CBT-I induce sex-specific responses and which domains are most affected (e.g., NREM oscillations, mood, memory, or subjective sleep quality), as evidence in this area remains incomplete and mixed. Understanding oscillatory signatures that differ by sex and age can guide targeted assessment and personalized interventions, moving the field toward mechanism-based, rather than symptom-based, treatment planning. Larger, well-powered, prospectively designed studies with uniformly characterized samples will be essential to refine these conclusions and clarify how age, sex, and insomnia jointly shape sleep physiology across the lifespan.

## Acknowledgments

We would like to thank all participants and volunteers for their work on these studies. A special thank you to our volunteer Isabella Di Matteo for her assistance with data entry and quality checking. We acknowledge the contributions of the following team members who assisted in participants’ recruitment, data collection and data preprocessing: Lukia Tarelli, Kirsten Gong, Margaret McCarthy, Ophelia Fontaine, Sam Gillman, Jean-Louis Zhao, and postdoctoral fellow Mathilde Reyt. We also thank our sleep technologists Madeline Dickson and Elinah Mozhentiy from the Clinique SomnoMed for their contribution to the setup of sleep recordings,

## Financial disclosure

TDV has received consultant and speaker fees from Eisai, Idorsia, Paladin Labs, Takeda and Axsome, as well as research grants from Jazz Pharmaceuticals and Paladin Labs.

## Non-financial disclosure

None to report.

## Funding

This work was funded by grants to TDV from the Natural Sciences and Engineering Research Council of Canada (NSERC), the Canadian Institutes of Health Research (MOP 142191, PJT153115) and the Fonds de Recherche du Québec (FRQ) and supported by a grant to NAW No. 767-2023-2413 from the Social Sciences and Humanities Research Council.

## Data availability

All code used in this study was developed by the authors for the execution of the study and is available online at https://github.com/nathanecross/seapipe. Raw data are restricted due to legal/ethical considerations. However, they may be shared with other investigators upon reasonable request and evaluation of such request by our local ethics review board

