## Supplemental Material for "Beyond Aging, Sex and Insomnia Disorder Shape NREM Brain Oscillations"

Supplemental Table 1 – *Age Differences in Sleep Macroarchitecture Variables*

|  | **Wald χ² (df= 1)** | **p** | **Coefficient** | **SE** | ***p*** | **95% Lower** | **Upper** | **Difference** |
| --- | --- | --- | --- | --- | --- | --- | --- | --- |
| **TIB (min)** | 0.56 | .445 | 0.05 | 0.23 | .807 | -.410 | .486 | - |
| **TST (min)** | 16.06 | < .001 | **-0.87** | **0.23** | **<.001** | **-1.34** | **-0.46** | **↓** |
| **SOL (min)** | 0.12 | .730 | 0.03 | 0.08 | .757 | -0.11 | 0.18 | - |
| **WASO (min)** | 33.89 | < .001 | **0.77** | **0.13** | **<.001** | **0.51** | **1.03** | **↑** |
| **SE (%)** | 41.53 | < .001 | **-0.22** | **0.04** | **<.001** | **-0.29** | **-0.16** | **↓** |
| **Wake (%)** | 37.71 | < .001 | **0.19** | **0.03** | **<.001** | **0.13** | **0.25** | **↑** |
| **NREM1 (%)** | 0.17 | .678 | 0.01 | 0.02 | .696 | -0.02 | 0.04 | - |
| **NREM2 (%)** | 11.78 | < .001 | **-0.39** | **0.11** | **.005** | **-0.62** | **-0.17** | **↓** |
| **NREM3 (%)** | 34.27 | < .001 | **-0.42** | **0.07** | **<.001** | **-0.57** | **-0.29** | **↓** |
| **REM (%)** | 16.99 | < .001 | **-0.27** | **0.07** | **.002** | **-0.40** | **-0.14** | **↓** |
| **SFI** | 42.24 | < .001 | **0.09** | **0.01** | **<.001** | **0.06** | **0.12** | **↑** |

Note. Bias-corrected confidence intervals are presented. *p* = adjusted p. Significant results are bolded. TIB = time in bed; TST = total sleep time; SOL = sleep onset latency; WASO = wake after sleep onset; SE= Sleep efficiency; NREM = non-rapid eye movement sleep; REM = rapid eye movement sleep; SFI = sleep fragmentation index;

Supplemental Table 2 – *Age Controlled Group Differences in Sleep Macroarchitecture Variables*

|  | **INS (n= 119)** | **HS (n= 103)** | **Wald χ² (df= 1)** | **p** | **Coefficient** | **SE** | ***p*** | **95% Lower** | | **Upper** | **Difference** |
| --- | --- | --- | --- | --- | --- | --- | --- | --- | --- | --- | --- |
| **TIB (min)** | 445.67 ± 5.31 | 453.51 ± 5.27 | 1.11 | .292 | -7.84 | 7.52 | .308 | -22.71 | | 6.35 | **-** |
| **TST (min)** | 364.51 ± 5.71 | 398.81 ± 4.84 | 20.96 | < .001 | **-34.30** | **7.53** | **<.001** | **-48.15** | | **-18.97** | **INS < HS** |
| **SOL (min)** | 14.89 ± 1.31 | 13.93 ± 1.54 | 0.21 | .645 | 0.96 | 2.14 | .650 | -3.78 | | 4.92 | - |
| **WASO (min)** | 58.80 ± 3.85 | 34.67 ± 2.15 | 27.80 | < .001 | **24.14** | **4.60** | **<.001** | **15.23** | | **32.94** | **INS > HS** |
| **SE (%)** | 81.85 ± 0.93 | 88.16 ± 0.61 | 30.21 | < .001 | **-6.30** | **1.14** | **<.001** | **-8.64** | **-4.09** | | **INS < HS** |
| **Wake (%)** | 13.83 ± 0.90 | 7.95 ± 0.47 | 31.39 | < .001 | **5.88** | **1.05** | **<.001** | **3.92** | **8.00** | | **INS > HS** |
| **NREM1 (%)** | 3.23 ± 0.21 | 5.03 ± 0.43 | 12.30 | <.001 | **-1.80** | **0.52** | **.007** | **-2.93** | **-0.82** | | **INS <HS** |
| **NREM2 (%)** | 44.41 ± 1.15 | 64.05 ± 4.95 | 18.72 | < .001 | **-19.65** | **4.61** | **.005** | **-29.11** | **-11.13** | | **INS < HS** |
| **NREM3 (%)** | 20.67 ± 0.87 | 31.93 ± 2.58 | 21.24 | < .001 | **-11.26** | **2.35** | **<.001** | **-16.36** | **-6.78** | | **INS < HS** |
| **REM (%)** | 22.90 ± 0.70 | 31.06 ± 2.58 | 11.57 | < .001 | **-8.16** | **2.42** | **.007** | **-13.07** | **-3.45** | | **INS < HS** |
| **SFI** | 7.66 ± 0.23 | 7.51 ± 0.27 | 0.16 | .686 | 0.15 | 0.37 | .676 | -0.59 | 0.85 | | - |

Note. Values are reported as estimated marginals means (age controlled) ± standard error with bias-corrected confidence intervals. *p* = adjusted p. Significant results are bolded. TIB = time in bed; TST = total sleep time; SOL = sleep onset latency; WASO = wake after sleep onset; SE= Sleep efficiency; NREM = non-rapid eye movement sleep; REM = rapid eye movement sleep; SFI = sleep fragmentation index; INS = insomnia group; HS = healthy sleeper group.

Supplemental Table 3 – *Age Controlled Sex Differences in Sleep Macroarchitecture Variables*

|  | **Male (n= 74)** | **Female (n= 148)** | **Wald χ² (df= 1)** | **p** | **Coefficient** | **SE** | ***p*** | **95% Lower** | | **Upper** | **Difference** |
| --- | --- | --- | --- | --- | --- | --- | --- | --- | --- | --- | --- |
| **TIB (min)** | 458.03 ± 6.30 | 444.95 ± 4.79 | 2.64 | .104 | 13.08 | 8.11 | .106 | -2.45 | | 29.22 | **-** |
| **TST (min)** | 383.28 ± 6.35 | 379.00 ± 5.09 | 0.27 | .604 | 4.28 | 8.14 | .601 | 11.66 | | 20.05 | **-** |
| **SOL (min)** | 13.31 ± 1.36 | 15.02 ± 1.24 | 0.90 | .343 | -1.70 | 1.80 | .348 | -5.40 | | 1.92 | **-** |
| **WASO (min)** | 53.99 ± 3.57 | 44.41 ± 2.91 | 4.49 | .034 | **9.58** | **4.65** | **.041** | **0.31** | | **18.57** | **M > F** |
| **SE (%)** | 83.73 ± 0.93 | 85.30 ± 0.74 | 1.78 | .182 | -1.57 | 1.15 | .173 | -3.82 | 0.77 | | **-** |
| **Wake (%)** | 12.39 ± 0.79 | 10.46 ± 0.69 | 3.51 | .061 | 1.93 | 1.05 | .059 | -0.10 | 4.01 | | **-** |
| **NREM1 (%)** | 5.15 ± 0.47 | 3.53 ± 0.21 | 9.94 | .002 | **1.62** | **0.49** | **.004** | **0.70** | **2.59** | | **M > F** |
| **NREM2 (%)** | 56.68 ± 5.35 | 51.94 ± 3.16 | 0.56 | .454 | 4.75 | 6.28 | .452 | -7.39 | 17.93 | | - |
| **NREM3 (%)** | 26.25 ± 3.14 | 25.72 ± 1.55 | 0.23 | .880 | 0.53 | 3.42 | .873 | -5.83 | 7.70 | | - |
| **REM (%)** | 27.65 ± 2.79 | 26.20 ± 1.65 | 0.19 | .661 | 1.45 | 3.28 | .659 | -4.64 | 8.01 | | - |
| **SFI** | 8.60 ± 0.24 | 7.09 ± 0.21 | 24.43 | < .001 | **1.52** | **0.31** | **< .001** | **0.93** | **2.13** | | **M > F** |

Note. Values are reported as estimated marginals means (age controlled) ± standard error with bias-corrected confidence intervals. *p* = adjusted p. Significant results are bolded. Sigma power (11.25-16Hz), SWA (0.25-4Hz), SO power (0.25-1.25 Hz), NREM2 + NREM3, M = male; F = female.

Supplemental Table 4- *Age Controlled* *GroupxSex Differences in Sleep Macroarchitecture Variables*

| **Variable** | **Male INS**  **(n= 34)** | **Female INS (n= 85)** | **Male HS**  **(n= 40)** | **Female HS**  **(n= 63)** | **Wald χ² (df)** | **p** | **Coefficient** | **SE** | ***p*** | **95% Lower** | **Upper** | **Difference** |
| --- | --- | --- | --- | --- | --- | --- | --- | --- | --- | --- | --- | --- |
| **Age (yrs)** | 41.29 ± 2.34 | 49.76 ± 1.58 | 34.00 ± 2.83 | 35.78 ± 2.32 | 38.79 (3) | <.001 | 8.47 | 2.82 | .005 | 2.33 | 14.25 | FINS > MINS |
|  |  |  |  |  |  |  | 13.99 | 2.72 | <.001 | 8.60 | 19.48 | FINS > FHS |
|  |  |  |  |  |  |  | -15.77 | 3.18 | <.001 | -22.05 | -9.23 | FINS > MHS |
|  |  |  |  |  |  |  | -7.29 | 3.67 | .039 | -14.30 | -0.06 | MINS > MHS |
| **TIB (min)** | 450.07 ± 9.95 | 443.63 ± 6.26 | 465.21 ± 7.70 | 446.47 ± 7.10 | 4.96 (3) | .175 | 21.58 | 10.32 | .028 | 3.19 | 40.49 | FINS < MHS |
| **TST (min)** | 360.71 ± 9.86 | 366.12 ± 7.07 | 404.92 ± 6.31 | 394.82 ± 6.81 | 25.10 (3) | <.001 | -44.21 | 11.68 | <.001 | -66.51 | -22.17 | MINS < MHS |
|  |  |  |  |  |  |  | -34.11 | 11.99 | .010 | -58.82 | -9.77 | MINS < FHS |
|  |  |  |  |  |  |  | 28.70 | 9.84 | .004 | 8.46 | 48.87 | FINS < FHS |
|  |  |  |  |  |  |  | -38.80 | 9.72 | <.001 | -57.03 | -20.34 | FINS < MHS |
| **SOL (min)** | 14.46 ± 1.96 | 15.10 ± 1.63 | 12.29 ± 1.94 | 14.93 ± 2.03 | 1.50 (3) | .682 | - | - | - | - | - | - |
| **WASO (min)** | 64.27 ± 6.49 | 56.32 ± 4.67 | 43.30 ± 2.69 | 29.56 ± 2.56 | 43.48 (3) | <.001 | -26.74 | 5.73 | <.001 | -38.99 | -15.57 | FINS > FHS |
|  |  |  |  |  |  |  | 13.03 | 5.78 | .032 | 2.49 | 24.25 | FINS > MHS |
|  |  |  |  |  |  |  | 20.97 | 7.30 | .008 | 6.93 | 36.98 | MINS > MHS |
|  |  |  |  |  |  |  | 34.68 | 6.91 | <.001 | 21.78 | 48.86 | MINS > FHS |
|  |  |  |  |  |  |  | -13.71 | 3.59 | <.001 | -20.30 | -7.14 | MHS > FHS |
| **SE (%)** | 80.18 ± 1.55 | 82.59 ± 1.15 | 87.23 ± .74 | 88.65 ± .81 | 33.83 (3) | <.001 | 7.05 | 1.73 | <.001 | 5.15 | 12.36 | MINS < MHS |
|  |  |  |  |  |  |  | 8.46 | 1.76 | <.001 | 5.15 | 12.36 | MINS < FHS |
|  |  |  |  |  |  |  | -6.05 | 1.40 | <.001 | -8.80 | -3.28 | FINS < FHS |
|  |  |  |  |  |  |  | -4.63 | 1.44 | .004 | -7.45 | -2.25 | FINS < MHS |
| **Wake (%)** | 15.07 ± 1.41 | 13.26 ± 1.12 | 9.63 ± .56 | 6.97 ± .59 | 44.30 (3) | <.001 | -6.29 | 1.34 | <.001 | -8.89 | -3.90 | FINS > FHS |
|  |  |  |  |  |  |  | -3.63 | 1.33 | .014 | -6.39 | -1.29 | FINS > MHS |
|  |  |  |  |  |  |  | 5.44 | 1.55 | .002 | 2.50 | 8.29 | MINS > MHS |
|  |  |  |  |  |  |  | 8.10 | 1.51 | <.001 | 4.82 | 11.25 | MINS > FHS |
|  |  |  |  |  |  |  | 2.66 | 0.75 | <.001 | 1.16 | 4.15 | MHS > FHS |
| **NREM1** | 3.84 ± .41 | 2.95 ± .24 | 6.37 ± .75 | 4.22 ± .44 | 19.59 (3) | <.001 | -3.42 | 0.88 | <.001 | -5.19 | -1.88 | FINS < MHS |
| **(%)** |  |  |  |  |  |  | -1.27 | 0.53 | .049 | -2.45 | -0.19 | FINS < FHS |
|  |  |  |  |  |  |  | 2.15 | 0.80 | .013 | 0.58 | 3.79 | FHS < MHS |
|  |  |  |  |  |  |  | 2.54 | 0.82 | .009 | 0.99 | 4.02 | MINS < MHS |
| **NREM2** | 43.38 ± 1.70 | 44.80 ± 1.48 | 69.38 ± 9.55 | 60.69 ± 6.05 | 19.47 (3) | <.001 | 15.89 | 5.68 | .022 | 5.97 | 27.83 | FINS < FHS |
| **(%)** |  |  |  |  |  |  | 24.57 | 9.41 | .025 | 8.80 | 43.16 | FINS < MHS |
|  |  |  |  |  |  |  | -26.00 | 10.12 | .037 | -47.06 | -7.38 | MINS < MHS |
|  |  |  |  |  |  |  | -15.89 | 5.67 | .020 | -27.82 | -5.17 | MINS < FHS |
| **NREM3 (%)** | 15.96 ± 1.18 | 22.70 ± 1.10 | 35.71 ± 5.52 | 29.33 ± 2.80 | 33.84 (3) | <.001 | 6.74 | 1.63 | <.001 | 3.58 | 9.92 | MINS < FINS |
|  |  |  |  |  |  |  | 19.75 | 5.89 | .009 | 9.24 | 31.83 | MINS < MHS |
|  |  |  |  |  |  |  | -13.37 | 3.11 | <.001 | -20.35 | -7.74 | MINS < FHS |
|  |  |  |  |  |  |  | -6.64 | 2.77 | .024 | -12.62 | -1.47 | FINS < FHS |
|  |  |  |  |  |  |  | -13.01 | 5.45 | .035 | -25.29 | -2.92 | FINS < MHS |
| **REM (%)** | 21.42 ± 1.29 | 23.52 ± .84 | 33.50 ± 4.96 | 29.46 ± 3.17 | 11.80 (3) | .008 | -8.04 | 3.38 | .042 | -14.94 | -1.67 | MINS < FHS |
|  |  |  |  |  |  |  | 12.07 | 5.20 | .035 | 3.05 | 22.89 | MINS < MHS |
|  |  |  |  |  |  |  | **-5.95** | **2.94** | **.057** | **-12.31** | **-0.32** | **FINS < FHS** |
|  |  |  |  |  |  |  | -9.98 | 5.01 | .07 | -21.55 | -1.21 | FINS < MHS |
| **SFI** | 8.77 ± .29 | 7.17 ± .30 | 8.45 ± .36 | 6.99 ± .32 | 27.13 (3) | <.001 | 1.78 | 0.45 | <.001 | 0.94 | 2.65 | MINS > FHS |
|  |  |  |  |  |  |  | -1.28 | 0.49 | .012 | -2.28 | -0.31 | FINS < MHS |
|  |  |  |  |  |  |  | 1.46 | 0.45 | <.001 | 0.61 | 2.32 | MHS > FHS |

Note. Values are reported as mean ± standard errors with bias-corrected confidence intervals. Only significant post-hoc analyses are presented. *p* = adjusted p; Results in bold are FDR-corrected comparisons p > .05 but became significant after bias-corrected confidence intervals. INS = insomnia group; HS = healthy sleeper group; TIB = time in bed; TST = total sleep time; SOL = sleep onset latency; WASO = wake after sleep onset; SE= Sleep efficiency; NREM = non-rapid eye movement sleep; REM = rapid eye movement sleep; SFI = sleep fragmentation index;

Supplemental Table 5- *Raw Means for Measures of Sleep Spindle and SWA (Fz)*

| **Variable (Fz)** | **INS**  **(n= 119)** | **HS**  **(n= 103)** | **Male**  **(n= 74)** | **Female**  **(n= 148)** | **Male INS**  **(n= 34)** | **Female INS**  **(n= 85)** | **Male HS**  **(n= 40)** | **Female HS**  **(n= 63)** |
| --- | --- | --- | --- | --- | --- | --- | --- | --- |
| **Event Detection** | | | | | | | | |
| Spindle Density | 2.07 ± .05 | 3.04 ± .084 | 2.54 ± .09 | 2.51 ± .71 | 2.16 ± .09 | 2.04 ± .05 | 2.87 ± .13 | 3.14 ± .11 |
| Spindle Amplitude | 104.50 ± 2.27 | 121.86 ± 4.34 | 106.46 ± 4.02 | 115.60 ± 3.00 | 99.04 ± 3.65 | 106.69 ± 2.80 | 112.78 ± 6.65 | 127.63 ± 5.62 |
| SO Density | 5.09 ± .09 | 7.92 ± .37 | 6.59 ± .38 | 6.31 ± .24 | 5.00 ± .19 | 5.13 ± .10 | 7.94 ± .62 | 7.90 ± .47 |
| SO Amplitude | 143.02 ± 3.86 | 182.24 ± 7.10 | 150.83 ± 6.65 | 166.41 ± 5.13 | 133.02 ± 5.92 | 147.01 ± 4.81 | 165.97 ± 10.73 | 192.57 ± 9.23 |
| **Power Spectral Analysis** | | | | | | | | |
| Absolute Sigma Power | 12.40 ± .64 | 13.71 ± .94 | 11.70 ± .80 | 13.62 ± .72 | 12.50 ± 1.25 | 12.36 ± .75 | 10.97 ± 1.02 | 15.40 ± 1.34 |
| Absolute SWA | 538.81 ± 30.32 | 817.43 ± 57.33 | 558.92 ± 43.30 | 715.40 ± 42.20 | 442.87 ± 36.13 | 577.18 ± 39.26 | 665.37 ± 72.36 | 911.21 ± 79.28 |
| Absolute SO Power | 436.71 ± 27.34 | 673.09 ± 51.37 | 481.04 ± 49.65 | 573.13 ± 34.88 | 356.30 ± 32.61 | 468.87 ± 35.50 | 595.67 ± 86.83 | 720.84 ± 63.26 |
| Relative Sigma Power | .020 ± .001 | .029 ± .001 | .027 ± .001 | .023 ± .001 | .024 ± .002 | .019 ± .001 | .031 ± .001 | .028 ± .001 |
| Relative SWA | .78 ± .01 | .80 ± .01 | .79 ± .01 | .80 ± .01 | .78 ± .01 | .79 ± .01 | .79 ± .01 | .81 ± .01 |
| Relative SO Power | .62 ± .01 | .64 ± .01 | .63 ± .01 | .63 ± .01 | .61 ± .02 | .62 ± .01 | .65 ± .02 | .63 ± .01 |

Note. Values are reported as raw mean ± standard error. INS = insomnia group; HS = healthy sleeper group. Sigma power (11.25-16Hz), SWA (0.25-4Hz), SO power (0.25-1.25 Hz). NREM2 + NREM3

Supplemental Table 6- *Raw Means for Measures of Sleep Spindle and SWA (Cz)*

| **Variable (Cz)** | **INS**  **(n= 119)** | **HS**  **(n= 103)** | **Male**  **(n= 74)** | **Female**  **(n= 148)** | **Male INS**  **(n= 34)** | **Female INS**  **(n= 85)** | **Male HS**  **(n= 40)** | **Female HS**  **(n= 63)** |
| --- | --- | --- | --- | --- | --- | --- | --- | --- |
| **Event Detection** | | | | | | | | |
| Spindle Density | 2.24 ± .06 | 3.27 ± .09 | 2.80 ± .10 | 2.68 ± .08 | 2.38 ± .12 | 2.19 ± .07 | 3.15 ± .14 | 3.34 ± .12 |
| Spindle Amplitude | 97.10 ± 2.27 | 107.95 ± 2.77 | 91.81 ± 2.31 | 107.30 ± 2.34 | 88.47 ± 2.79 | 100.56 ± 2.90 | 94.65 ± 3.52 | 116.39 ± 3.57 |
| SO Density | 5.00 ± .09 | 7.83 ± .38 | 6.48 ± .39 | 6.22 ± .24 | 4.95 ± .19 | 5.01 ± .11 | 7.78 ± .63 | 7.86 ± .49 |
| SO Amplitude | 129.90 ± 3.18 | 155.83 ± 4.67 | 129.05 ± 4.00 | 148.36 ± 3.74 | 119.14 ± 4.58 | 134.20 ± 3.97 | 137.48 ± 6.02 | 167.47 ± 6.21 |
| **Power Spectral Analysis** | | | | | | | | |
| Absolute Sigma Power | 15. 69 ± .78 | 17.05 ± 1.00 | 14.28 ± .95 | 17.29 ± .79 | 14.98 ± 1.41 | 15.98 ± .94 | 13.64 ± 1.28 | 19.16 ± 1.35 |
| Absolute SWA | 417.01 ± 22.43 | 647.44 ± 42.75 | 440.29 ± 35.44 | 559.76 ± 30.95 | 340.89 ± 25.77 | 447.46 ± 29.10 | 531.64 ± 60.36 | 718.86 ± 56.67 |
| Absolute SO Power | 324.76 ± 20.48 | 543.66 ± 48.39 | 390.25 ± 55.15 | 439.12 ± 26.83 | 261.29 ± 23.57 | 350.15 ± 26.66 | 508.76 ± 100.32 | 565.17 ± 48.50 |
| Relative Sigma Power | .031 ± .001 | .025 ± .001 | .029 ± .002 | .028 ± .001 | .034 ± .002 | .029 ± .001 | .025 ± .002 | .026 ± .002 |
| Relative SWA | .75 ± .01 | .77 ± .01 | .75 ± .01 | .76 ± .01 | .75 ± .01 | .75 ± .01 | .76 ± .013 | .78 ± .01 |
| Relative SO Power | .56 ± .01 | .60 ± .01 | .58 ± .01 | .58 ± .01 | .56 ± .02 | .56 ± .01 | .61 ± .02 | .59 ± .01 |

Note. Values are reported as raw mean ± standard error. INS = insomnia group; HS = healthy sleeper group. Sigma power (11.25-16Hz), SWA (0.25-4Hz), SO power (0.25-1.25 Hz). NREM2 + NREM3

Supplemental Table 7- *Raw Means for Measures of Sleep Spindle and SWA (Pz)*

| **Variable (Pz)** | **INS**  **(n= 119)** | **HS**  **(n= 103)** | **Male**  **(n= 74)** | **Female**  **(n= 148)** | **Male INS**  **(n= 34)** | **Female INS**  **(n= 85)** | **Male HS**  **(n= 40)** | **Female HS**  **(n= 63)** |
| --- | --- | --- | --- | --- | --- | --- | --- | --- |
| **Event Detection** | | | | | | | | |
| Spindle Density | 2.46 ± .62 | 3.58 ± .11 | 3.06 ± .12 | 2.93 ± .09 | 2.60 ± .11 | 2.40 ± .07 | 3.45 ± .17 | 3.66 ± .14 |
| Spindle Amplitude | 79.46 ± 1.75 | 96.32 ± 3.88 | 80.92 ± 4.13 | 90.46 ± 2.35 | 72.30 ± 2.30 | 82.33 ± 2.21 | 88.26 ± 7.23 | 101.44 ± 4.31 |
| SO Density | 4.71 ± .09 | 7.35 ± .36 | 6.06 ± .37 | 5.87 ± .23 | 4.59 ± .19 | 4.76 ± .10 | 7.31 ± .60 | 7.37 ± .46 |
| SO Amplitude | 110.51 ± 2.83 | 139.49 ± 5.63 | 112.65 ± 5.40 | 129.61± 3.84 | 100.57 ± 4.18 | 114.48 ± 3.52 | 122.91 ± 9.07 | 150.01 ± 6.92 |
| **Power Spectral Analysis** | | | | | | | | |
| Absolute Sigma Power | 13.98 ± .77 | 14.66 ± .87 | 12.41 ± .90 | 15.20 ± .72 | 12.71 ± 1.17 | 14.49 ± .97 | 12.14 ± 1.37 | 16.21 ± 1.09 |
| Absolute SWA | 318.11 ± 18.08 | 467.81 ± 33.90 | 309.71 ± 26.07 | 422.36 ± 24.47 | 247.00 ± 18.96 | 346.55 ± 23.51 | 367.34 ± 45.18 | 529.76 ± 45.66 |
| Absolute SO Power | 278.07 ± 18.91 | 431.12 ± 39.57 | 303.06 ± 43.43 | 368.22 ± 23.24 | 213.60 ± 20.52 | 303.86 ± 24.69 | 385.27 ± 79.32 | 459.39 ± 41.42 |
| Relative Sigma Power | .036 ± .001 | .038 ± .001 | .041 ± .002 | .035 ± .001 | .040 ± .003 | .034 ± .002 | .042 ± .002 | .036 ± .002 |
| Relative SWA | .72 ± .01 | .74 ± .01 | .72 ± .01 | .74 ± .01 | .72 ± .01 | .72 ± .01 | .73 ± .01 | .75 ± .01 |
| Relative SO Power | .60 ± .01 | .64 ± .01 | .62 ± .01 | .62 ± .01 | .59 ± .02 | .61 ± .01 | .65 ± .02 | .63 ± .01 |

Note. Values are reported as raw mean ± standard error. INS = insomnia group; HS = healthy sleeper group. Sigma power (11.25-16Hz), SWA (0.25-4Hz), SO power (0.25-1.25 Hz). NREM2 + NREM3

Supplemental Table 8- *Age Differences in Sigma and SWA Across All Participants*

| **Variable** | **Wald χ² (df= 1)** | **p** | **Coefficient** | **SE** | ***p*** | **95% Lower** | **Upper** | **Difference** |
| --- | --- | --- | --- | --- | --- | --- | --- | --- |
| **Fz** | | | | | | | | |
| **Event Detection** | | | | | | | | |
| Spindle Amplitude | 82.37 | <.001 | **-0.99** | **0.107** | **< .001** | **-1.20** | **-0.78** | **↓** |
| SO Amplitude | 122.93 | .000 | **-1.95** | **0.185** | **< .001** | **-2.35** | **-1.58** | **↓** |
| **Power Spectral Analysis** | | | | | | | | |
| Relative SO power | 27.26 | < .001 | **-0.002** | **0.001** | **<.001** | **-0.003** | **-0.001** | **↓** |
| Absolute Sigma power | 47.48 | < .001 | **-0.195** | **.029** | **<.001** | **-0.253** | **-0.138** | **↓** |
| Absolute SWA | 92.28 | < .001 | **-14.33** | **1.54** | **< .001** | **-17.57** | **-11.24** | **↓** |
| Absolute SO power | 70.89 | < .001 | **-12.00** | **1.43** | **< .001** | **-14.97** | **-9.05** | **↓** |
| **Cz** | | | | | | | | |
| **Event Detection** | | | | | | | | |
| Spindle Amplitude | 88.35 | .000 | **-0.76** | **0.08** | **< .001** | **-0.92** | **-0.62** | **↓** |
| SO Amplitude | 131.24 | .000 | **-1.44** | **0.13** | **< .001** | **-1.70** | **-1.19** | **↓** |
| **Power Spectral Analysis** | | | | | | | | |
| Relative SO power | 36.10 | < .001 | **-0.003** | **0.000** | **<.001** | **-0.004** | **-0.002** | **↓** |
| Absolute Sigma power | 42.18 | < .001 | **-0.206** | **.033** | **< .001** | **-0.270** | **-0.139** | **↓** |
| Absolute SWA | 98.75 | < .001 | **-10.81** | **1.09** | **< .001** | **-13.06** | **-8.60** | **↓** |
| Absolute SO power | 65.35 | < .001 | **-9.64** | **1.17** | **< .001** | **-11.82** | **-7.49** | **↓** |
| **Pz** | | | | | | | | |
| **Event Detection** | | | | | | | | |
| Spindle Density | 10.31 | < .001 | **-0.01** | **0.004** | **.003** | **-0.02** | **-0.004** | **↓** |
| Spindle Amplitude | 63.12 | <.001 | **-0.71** | **0.089** | **< .001** | **-0.91** | **-0.56** | **↓** |
| SO Density | 2.15 | .142 | -0.02 | 0.011 | .160 | -0.037 | 0.006 | - |
| SO Amplitude | 109.73 | .000 | **-1.40** | **0.136** | **< .001** | **-1.69** | **-1.15** | **↓** |
| **Power Spectral Analysis** | | | | | | | | |
| Relative Sigma Power | 2.38 | .123 | 8.65 × 10^-5^ | 5.71 × 10^-5^ | .126 | -2.29 × 10^-5^ | 0.000 | - |
| Relative SWA | 39.29 | < .001 | **-0.002** | **.000** | **< .001** | **-0.002** | **-0.001** | **↓** |
| Relative SO power | 30.35 | < .001 | **-0.002** | **.000** | **<.001** | **-0.003** | **-0.002** | **↓** |
| Absolute Sigma power | 23.39 | <.001 | **-0.141** | **.029** | **<.001** | **-0.203** | **-0.077** | **↓** |
| Absolute SWA | 69.69 | < .001 | **-7.49** | **.926** | **< .001** | **-9.22** | **-5.89** | **↓** |
| Absolute SO power | 46.71 | < .001 | **-7.01** | **1.05** | **< .001** | **-9.28** | **-4.91** | **↓** |

Note. Bias-corrected confidence intervals are presented. *p* = adjusted p; Significant results are bolded. Sigma power (11.25-16Hz), SWA (0.25-4Hz), SO power (0.25-1.25 Hz). NREM2 + NREM3

Supplemental Table 9- *Age Controlled Group Differences in Measures of Spindle and SWA*

| **Variable** | **INS (n= 119)** | **HS (n= 103)** | **Wald χ² (df= 1)** | **p** | **Coefficient** | **SE** | **p** | **95% Lower** | **Upper** | **Difference** |
| --- | --- | --- | --- | --- | --- | --- | --- | --- | --- | --- |
| **Fz** | | | | | | | | | | |
| **Event Detection** | | | | | | | | | | |
| Spindle Amplitude | 106.65 ± 2.30 | 113.52 ± 3.33 | 3.17 | .075 | -6.87 | 3.69 | .07 | -14.27 | 0.55 | - |
| SO Amplitude | 147.60 ± 3.58 | 166.07 ± 5.13 | 9.13 | .003 | **–18.47** | **6.15** | **.004** | **–31.06** | **–7.08** | **INS < HS** |
| **Power Spectral Analysis** | | | | | | | | | | |
| Relative SO Power | .63 ± .01 | .62 ± .01 | .024 | .877 | 0.002 | 0.012 | .892 | –0.022 | 0.026 | - |
| Absolute Sigma Power | 13.34 ± .65 | 12.56 ± .77 | 0.41 | .520 | 0.63 | 0.99 | .515 | -1.27 | 2.55 | - |
| Absolute SWA | 607.05 ± 30.95 | 733.72 ± 43.96 | 7.81 | .005 | **-139.57** | **51.78** | **.009** | **-243.51** | **-40.99** | **INS < HS** |
| Absolute SO Power | 494.66 ± 28.21 | 601.99 ± 40.85 | 6.15 | .013 | **–116.20** | **47.48** | **.015** | **–215.05** | **-25.60** | **INS < HS** |
| **Cz** | | | | | | | | | | |
| **Event Detection** | | | | | | | | | | |
| Spindle Amplitude | 96.97 ± 2.05 | 100.63 ± 1.90 | 1.60 | .206 | –3.66 | 2.91 | .202 | –9.59 | 2.13 | - |
| SO Amplitude | 131.68 ± 2.78 | 143.01 ± 3.14 | 6.72 | .010 | **-11.33** | **4.33** | **.007** | **-19.28** | **-2.91** | **INS < HS** |
| **Power Spectral Analysis** | | | | | | | | | | |
| Relative SO Power | .57 ± .01 | .58 ± .01 | 0.22 | .638 | -0.007 | 0.014 | .523 | –0.038 | 0.020 | - |
| Absolute Sigma Power | 16.67 ± .78 | 15.86 ± .85 | 0.29 | .591 | 0.60 | 1.11 | .582 | –1.57 | 2.73 | - |
| Absolute SWA | 468.41 ± 23.18 | 584.39 ± 32.50 | 11.68 | < .001 | **–125.89** | **38.12** | **.004** | **–195.50** | **–45.01** | **INS < HS** |
| Absolute SO Power | 371.79 ± 21.58 | 485.96 ± 38.61 | 9.03 | .003 | **-120.20** | **40.20** | **.007** | **-201.50** | **-41.25** | **INS < HS** |
| **Pz** | | | | | | | | | | |
| **Event Detection** | | | | | | | | | | |
| Spindle Density | 2.51 ± .07 | 3.50 ± .11 | 57.14 | <.001 | **–0.99** | **0.127** | **<.001** | **–1.23** | **-0.72** | **INS < HS** |
| Spindle Amplitude | 80.25 ± 1.79 | 89.93 ± 3.23 | 9.28 | .002 | **-9.68** | **3.25** | **.006** | **-16.04** | **-3.57** | **INS < HS** |
| SO Density | 4.76 ± .13 | 7.22 ± .35 | 45.59 | <.001 | **-2.46** | **0.382** | **<.001** | **-3.22** | **-1.74** | **INS < HS** |
| SO Amplitude | 112.54 ± 2.66 | 127.18 ± 4.20 | 10.00 | .002 | **-14.64** | **4.72** | **.002** | **-24.00** | **-5.83** | **INS < HS** |
| **Power Spectral Analysis** | | | | | | | | | | |
| Relative Sigma Power | .04 ± .002 | .04 ± .001 | 2.02 | .155 | -0.03 | 0.002 | .148 | –0.007 | 0.001 | - |
| Relative SWA | .73 ± .01 | .73 ± .01 | 0.01 | .920 | –0.001 | 0.009 | .926 | –0.019 | 0.017 | - |
| Relative SO Power | .61 ± .01 | .62 ± .01 | 0.23 | .632 | -0.007 | 0.014 | .634 | -0.033 | 0.021 | - |
| Absolute Sigma Power | 14.63 ± .78 | 13.86 ± .79 | 0.31 | .580 | 0.60 | 1.09 | .586 | –1.61 | 2.64 | - |
| Absolute SWA | 353.14 ± 18.71 | 424.83 ± 26.84 | 7.23 | .007 | **–79.99** | **29.84** | **.007** | **–138.21** | **–18.24** | **INS < HS** |
| Absolute SO Power | 311.64 ± 19.63 | 389.94 ± 32.51 | 6.04 | .014 | **-84.14** | **35.69** | **.020** | **-161.03** | **-18.82** | **INS < HS** |

Note. Values are reported as estimated marginals means (age controlled) ± standard error with bias-corrected confidence intervals. *p* = adjusted p; Significant results are bolded. Sigma power (11.25-16Hz), SWA (0.25-4Hz), SO power (0.25-1.25 Hz). INS = insomnia group; HS = healthy sleeper group. NREM2 + NREM3

Supplemental Table 10- *Age Controlled Sex Differences in Measures of Spindle and SWA*

| **Variable** | **Male (n= 74)** | **Female (n= 148)** | **Wald χ² (df= 1)** | **p** | **Coefficient** | **SE** | ***p*** | **95% Lower** | **Upper** | **Difference** |
| --- | --- | --- | --- | --- | --- | --- | --- | --- | --- | --- |
| **Fz** | | | | | | | | | | |
| **Event Detection** | | | | | | | | | | |
| Spindle Amplitude | 101.91 ± 3.35 | 118.25 ± 2.73 | 13.74 | <.001 | **-16.34** | **4.62** | **<.001** | **-25.34** | **-7.25** | **M < F** |
| SO Amplitude | 141.70 ± 4.94 | 171/97 ± 4.32 | 20.28 | <.001 | **–30.27** | **6.63** | **<.001** | **–42.40** | **-16.65** | **M < F** |
| **Power Spectral Analysis** | | | | | | | | | | |
| Relative SO power | .62 ± .01 | .63 ± .01 | 0.44 | .508 | -0.008 | .012 | .519 | -0.031 | 0.016 | - |
| Absolute Sigma power | 10.91 ± .70 | 14.01 ± .69 | 8.71 | .003 | **-3.07** | **1.04** | **.004** | **-5.24** | **-0.98** | **M < F** |
| Absolute SWA | 493.51 ± 33.21 | 747.38 ± 36.62 | 25.69 | <.001 | **–260.99** | **52.16** | **<.001** | **–359.53** | **-160.73** | **M < F** |
| Absolute SO power | 426.37 ± 40.23 | 599.90 ± 30.57 | 12.58 | < .001 | **–179.46** | **50.80** | **.002** | **–268.24** | **-84.09** | **M < F** |
| **Cz** | | | | | | | | | | |
| **Event Detection** | | | | | | | | | | |
| Spindle Amplitude | 88.39 ± 1.90 | 109.21 ± 1.98 | 55.33 | <.001 | **–20.82** | **2.87** | **<.001** | **–26.56** | **–15.08** | **M < F** |
| SO Amplitude | 122.39 ± 2.91 | 152.31 ± 2.98 | 48.27 | <.001 | **–29.92** | **4.34** | **<.001** | **–38.51** | **–21.28** | **M < F** |
| **Power Spectral Analysis** | | | | | | | | | | |
| Relative SO power | .57 ± .01 | .58 ± .01 | 0.44 | .507 | -0.009 | .014 | .523 | –0.038 | 0.020 | - |
| Absolute Sigma power | 13.44 ± .86 | 17.71 ± .75 | 12.85 | <.001 | **–4.24** | **1.23** | **.002** | **–6.70** | **–1.89** | **M < F** |
| Absolute SWA | 390.25 ± 27.73 | 584.26 ± 26.20 | 27.02 | <.001 | **–200.44** | **39.14** | **<.001** | **–277.40** | **-125.47** | **M < F** |
| Absolute SO power | 345.31 ± 46.46 | 461.13 ± 23.23 | 5.99 | .014 | **-121.95** | **50.67** | **.039** | **-216.14** | **-14.27** | **M < F** |
| **Pz** | | | | | | | | | | |
| **Event Detection** | | | | | | | | | | |
| Spindle Density | 2.97 ± .10 | 3.03 ± .08 | 0.28 | .598 | -0.06 | .128 | .610 | –0.33 | 0.16 | - |
| Spindle Amplitude | 77.46 ± 3.47 | 92.72 ± 2.20 | 14.07 | <.001 | **-15.26** | **4.15** | **.002** | **-22.81** | **-6.00** | **M < F** |
| SO Density | 5.89 ± .32 | 6.09 ± .22 | 0.25 | .617 | –0.197 | .406 | .634 | –0.985 | 0.593 | - |
| SO Amplitude | 106.02 ± 4.20 | 133.71 ± 3.27 | 26.62 | <.001 | **-27.70** | **5.43** | **<.001** | **-38.27** | **-17.28** | **M < F** |
| **Power Spectral Analysis** | | | | | | | | | | |
| Relative Sigma power | .04 ± .002 | .04 ± .001 | 7.79 | .005 | **0.006** | **.002** | **.008** | **0.002** | **0.011** | **M > F** |
| Relative SWA | .72 ± .01 | .74 ± .01 | 5.98 | .014 | **–0.024** | **.010** | **.016** | **–0.044** | **–0.006** | **M < F** |
| Relative SO power | .61 ± .01 | .62 ± .01 | 0.62 | .430 | -0.011 | .014 | .451 | -0.040 | 0.017 | - |
| Absolute Sigma power | 11.84 ± .80 | 15.48 ± .72 | 11.24 | <.001 | **-3.61** | **1.11** | **.002** | **-5.90** | **-1.32** | **M < F** |
| Absolute SWA | 275.29 ± 20.76 | 439.22 ± 22.05 | 29.16 | <.001 | **-168.01** | **31.25** | **<.001** | **–230.61** | **-104.35** | **M < F** |
| Absolute SO power | 270.53 ± 37.07 | 384.15 ± 21.29 | 8.15 | .004 | **-117.91** | **42.64** | **.018** | **-191.95** | **-33.29** | **M < F** |

Note. Values are reported as estimated marginals means (age controlled) ± standard error with bias-corrected confidence intervals. *p* = adjusted p; Significant results are bolded. Sigma power (11.25-16Hz), SWA (0.25-4Hz), SO power (0.25-1.25 Hz). M = male; F = female. NREM2 + NREM3

Supplemental Table 11- *Age Controlled Group by Sex Differences in Spindle and SWA Measures (Fz)*

| **(Fz)** | **Male INS**  **(n= 34)** | **Female INS**  **(n= 85)** | **Male HS**  **(n= 40)** | **Female HS**  **(n= 63)** | **Wald χ² (df= 3)** | **p** | **Coefficient** | **SE** | ***p*** | **95% Lower** | **Upper** | **Difference** |
| --- | --- | --- | --- | --- | --- | --- | --- | --- | --- | --- | --- | --- |
| **Event Detection** | | | | | | | | | | | | |
| Spindle | 98.68 ± 3.30 | 114.73 ± 2.90 | 105.18 ± 5.55 | 121.80 ± 4.51 | 20.94 | <.001 | -23.13 | 5.72 | <.001 | -33.69 | -11.43 | MINS < FHS |
| Amplitude |  |  |  |  |  |  | -16.06 | 4.49 | <.001 | -24.61 | -7.29 | MINS < FINS |
|  |  |  |  |  |  |  | 16.62 | 7.49 | .033 | 2.39 | 30.64 | MHS < FHS |
| SO | 132.31 ± 4.77 | 162.80 ± 4.70 | 151.06 ± 8.25 | 181.12 ± 7.13 | 37.45 | <.001 | 48.80 | 8.73 | <.001 | 32.40 | 67.06 | MINS < FHS |
| Amplitude |  |  |  |  |  |  | -18.32 | 8.41 | .021 | –36.04 | –2.99 | FINS < FHS |
|  |  |  |  |  |  |  | 30.49 | 6.89 | <.001 | 17.62 | 44.29 | MINS < FINS |
|  |  |  |  |  |  |  | -30.06 | 11.28 | .014 | -52.41 | -7.47 | MHS < FHS |
|  |  |  |  |  |  |  | **18.74** | **9.63** | **.054** | **0.53** | **38.50** | **MINS < MHS** |
| **Power Spectral Analysis** | | | | | | | | | | | | |
| Relative SO | .61 ± .02 | .64 ± .01 | .63 ± .01 | .62 ± .012 | 3.01 | .390 | - | - | - | - | - | - |
| Power |  |  |  |  |  |  |  |  |  |  |  |  |
| Absolute Sigma | 12.32 ± 1.08 | 13.80 ± .78 | 9.60 ± .90 | 14.30 ± 1.12 | 13.87 | .003 | -4.71 | 1.53 | .004 | -8.09 | -1.51 | MHS < FHS |
| Power |  |  |  |  |  |  | **2.73** | **1.41** | **.053** | **0.19** | **5.48** | **MINS > MHS** |
|  |  |  |  |  |  |  | 4.21 | 1.26 | <.001 | 2.01 | 7.30 | MHS < FINS |
| Absolute | 429.86 ± 32.65 | 685.41 ± 39.27 | 562.26 ± 53.92 | 828.85 ± 62.32 | 41.70 | <.001 | -399.00 | 74.29 | <.001 | -573.47 | -260.92 | MINS < FHS |
| SWA |  |  |  |  |  |  | -255.55 | 55.49 | <.001 | –374.94 | –137.67 | MINS < FINS |
| Power |  |  |  |  |  |  | 266.59 | 88.04 | <.001 | 108.08 | 432.41 | MHS < FHS |
|  |  |  |  |  |  |  | **143.45** | **74.28** | **.053** | **15.98** | **287.34** | **FINS < FHS** |
|  |  |  |  |  |  |  | -132.41 | 63.23 | .049 | -276.80 | -14.46 | MINS < MHS |
| Absolute | 345.29 ± 32.48 | 560.38 ± 36.07 | 508.48 ± 69.55 | 651.20 ± 51.91 | 32.78 | <.001 | 305.91 | 60.30 | <.001 | 201.20 | 432.44 | MINS < FHS |
| SO Power |  |  |  |  |  |  | 215.09 | 51.86 | <.001 | 115.36 | 329.78 | MINS < FINS |
|  |  |  |  |  |  |  | **163.19** | **75.94** | **.073** | **30.35** | **305.60** | **MINS < MHS** |

Note. Values are reported as estimated marginals means (age controlled) ± standard error with bias-corrected confidence intervals. Only significant post-hoc analyses are presented. *p* = adjusted p; Results in bold are FDR-corrected comparisons p > .05 but became significant after bias-corrected confidence intervals. Sigma power (11.25-16Hz), SWA (0.25-4Hz), SO power (0.25-1.25Hz). NREM2 + NREM3

Supplemental Table 12- *Age Controlled Group by Sex Differences in Spindle and SWA Measures (Cz)*

| **Cz** | **Male INS**  **(n= 34)** | **Female INS**  **(n= 85)** | **Male HS**  **(n= 40)** | **Female HS**  **(n= 63)** | ***Wald χ²* (df= 3)** | **p** | **Coefficient** | **SE** | ***p*** | **95% Lower** | **Upper** | **Difference** |
| --- | --- | --- | --- | --- | --- | --- | --- | --- | --- | --- | --- | --- |
| **Event Detection** | | | | | | | | | | | | |
| Spindle | 88.19 ± 2.55 | 106.67 ± 2.89 | 88.88 ± 2.77 | 111.96 ± 2.70 | 59.13 | <.001 | 23.76 | 3.81 | <.001 | 16.15 | 31.28 | MINS < FHS |
| Amplitude |  |  |  |  |  |  | –23.08 | 3.98 | <.001 | –31.26 | -15.47 | MHS < FHS |
|  |  |  |  |  |  |  | 18.48 | 3.88 | <.001 | 11.02 | 25.95 | MINS < FINS |
|  |  |  |  |  |  |  | 17.80 | 4.28 | <.001 | 9.37 | 25.95 | FINS > MHS |
| SO | 118.62 ± 3.66 | 145.83 ± 3.81 | 126.50 ± 4.43 | 159.04 ± 4.59 | 56.91 | <.001 | –40.42 | 5.95 | <.001 | 29.64 | 53.15 | MINS < FHS |
| Amplitude |  |  |  |  |  |  | –32.53 | 6.74 | <.001 | –45.78 | –19.01 | MHS < FHS |
|  |  |  |  |  |  |  | 27.21 | 5.22 | <.001 | 17.62 | 38.29 | MINS < FINS |
|  |  |  |  |  |  |  | 19.33 | 6.20 | .002 | 6.92 | 31.50 | FINS > MHS |
|  |  |  |  |  |  |  | -13.21 | 5.96 | .031 | -24.56 | -1.71 | FINS < FHS |
| **Power Spectral Analysis** | | | | | | | | | | | | |
| Relative SO Power | .55 ± .02 | .59 ± .01 | .59 ± .02 | .57 ± .01 | 3.82 | .281 | - | - | - | - | - | - |
| Absolute Sigma | 14.80 ± 1.29 | 17.50 ± .94 | 12.18 ± 1.14 | 18.00 ± 1.15 | 16.25 | .001 | 5.81 | 1.64 | <.001 | 2.88 | 8.92 | MHS < FHS |
| Power |  |  |  |  |  |  | -5.32 | 1.51 | .002 | -8.13 | -2.50 | MHS < FINS |
| Absolute | 331.07 ± 23.70 | 529.13 ± 29.46 | 453.83 ± 46.54 | 656.71 ± 43.40 | 53.01 | <.001 | 325.64 | 48.33 | <.001 | 232.59 | 433.77 | MINS < FHS |
| SWA |  |  |  |  |  |  | –127.58 | 52.15 | .016 | –224.17 | –30.01 | FINS < FHS |
| Power |  |  |  |  |  |  | 198.06 | 38.96 | <.001 | 126.56 | 280.86 | MINS < FINS |
|  |  |  |  |  |  |  | 122.76 | 52.28 | .036 | 28.66 | 224.89 | MINS < MHS |
|  |  |  |  |  |  |  | -202.88 | 67.45 | .005 | -322.39 | -80.38 | MHS < FHS |
| Absolute | 252.38 ± 23.23 | 424.14 ± 27.75 | 438.27 ± 85.57 | 508.86 ± 39.33 | 41.69 | <.001 | –256.48 | 45.63 | <.001 | –354.17 | -167.74 | MINS < FHS |
| SO Power |  |  |  |  |  |  | -171.78 | 38.83 | <.001 | -245.92 | -105.62 | MINS < FINS |
|  |  |  |  |  |  |  | **-185.88** | **89.00** | **.114** | **-399.30** | **-18.72** | **MINS < MHS** |

Note. Values are reported as estimated marginals means (age controlled) ± standard error with bias-corrected confidence intervals. Only significant post-hoc analyses are presented. *p* = adjusted p; Results in bold are FDR-corrected comparisons p > .05 but became significant after bias-corrected confidence intervals. Sigma power (11.25-16Hz), SWA (0.25-4Hz), SO power (0.25-1.25Hz). NREM2 + NREM3

Supplemental Table 13- *Age Controlled Group by Sex Differences in Spindle and SWA Measures (Pz)*

| **(Pz)** | **Male INS**  **(n= 34)** | **Female INS**  **(n= 85)** | **Male HS**  **(n= 40)** | **Female HS**  **(n= 63)** | ***Wald χ²* (df= 3)** | **p** | **Coefficient** | **SE** | ***p*** | **95% Lower** | **Upper** | **Difference** |
| --- | --- | --- | --- | --- | --- | --- | --- | --- | --- | --- | --- | --- |
| **Event Detection** | | | | | | | | | | | | |
| Spindle  Density | 2.59 ± .10 | 2.49 ± .08 | 3.37 ± .16 | 3.60 ± .15 | 58.99 | <.001 | 0.77 | 0.19 | <.001 | 0.384 | 1.151 | MINS < MHS |
|  |  |  |  |  |  |  | 1.00 | 0.17 | <.001 | 0.697 | 1.359 | MINS < FHS |
|  |  |  |  |  |  |  | -0.88 | 0.18 | <.001 | -1.228 | -0.526 | FINS < MHS |
|  |  |  |  |  |  |  | -1.11 | 0.17 | <.001 | -1.45 | -0.792 | FINS < FHS |
| Spindle | 72.04 ± 1.84 | 88.13 ± 2.34 | 82.77 ± 6.28 | 97.23 ± 3.54 | 50.71 | <.001 | 25.20 | 4.04 | <.001 | 17.85 | 33.71 | MINS < FHS |
| Amplitude |  |  |  |  |  |  | -9.11 | 4.00 | .028 | -17.17 | -1.34 | FINS < FHS |
|  |  |  |  |  |  |  | 16.09 | 3.06 | <.001 | 10.30 | 22.27 | MINS < FINS |
| SO Density | 4.58 ± .19 | 4.89 ± .14 | 7.19 ± .58 | 7.28 ± .44 | 46.97 | <.001 | 2.39 | .45 | <.001 | 1.458 | 3.255 | FINS < FHS |
|  |  |  |  |  |  |  | -2.60 | .61 | .002 | -3.89 | -1.48 | MINS < MHS |
|  |  |  |  |  |  |  | -2.70 | .50 | <.001 | -3.72 | -1.72 | MINS < FHS |
|  |  |  |  |  |  |  | 2.30 | .62 | .004 | 1.076 | 3.538 | FINS < MHS |
| SO | 100.06 ± 3.09 | 125.80 ± 3.55 | 112.22 ± 7.37 | 141.80 ± 5.35 | 55.14 |  | –41.74 | 6.23 | <.001 | –55.33 | –30.97 | MINS < FHS |
| Amplitude |  |  |  |  |  |  | 16.00 | 6.23 | .012 | 3.82 | 28.85 | FINS < FHS |
|  |  |  |  |  |  |  | 29.58 | 9.64 | .005 | 9.92 | 47.84 | MHS < FHS |
|  |  |  |  |  |  |  | -25.74 | 4.84 | <.001 | -35.74 | -16.43 | MINS < FINS |
| **Power Spectral Analysis** | | | | | | | | | | | | |
| Relative Sigma  Power | .04 ± .003 | .03 ± .002 | .04 ± .002 | .04 ± .002 | 12.47 | .006 | 0.006 | 0.00 | .025 | 0.001 | 0.011 | MHS > FHS |
|  |  |  |  |  |  |  | -0.009 | 0.00 | .003 | -0.015 | -0.003 | FINS < MHS |
| Relative SWA | .72 ± .01 | .74 ± .01 | .71 ± .013 | .74 ± .01 | 6.17 | .104 | - | - | - | - | - | - |
| Relative SO Power | .59 ± .02 | .63 ± .01 | .63 ± .02 | .62 ± .01 | 3.82 | .282 | - | - | - | - | - | - |
| Absolute | 12.58 ± 1.04 | 15.54 ± .99 | 11.13 ± 1.22 | 15.41 ± 1.02 | 11.60 | .009 | -4.27 | 1.67 | .011 | -7.57 | -1.02 | MHS < FHS |
| Sigma |  |  |  |  |  |  | 2.96 | 1.39 | .040 | 0.41 | 5.57 | MINS < FINS |
| Power |  |  |  |  |  |  | 4.41 | 1.60 | .014 | 1.67 | 7.57 | MHS < FINS |
| Absolute | 240.20 ± 15.57 | 403.10 ± 24.31 | 313.46 ± 36.26 | 486.73 ± 36.72 | 54.12 | <.001 | –246.53 | 41.23 | <.001 | -330.71 | -170.69 | MINS < FHS |
| SWA |  |  |  |  |  |  | **83.63** | **43.14** | **.061** | **2.52** | **161.16** | **FINS < FHS** |
| Power |  |  |  |  |  |  | -162.90 | 30.05 | <.001 | -226.65 | -110.70 | MINS < FINS |
|  |  |  |  |  |  |  | -89.65 | 42.00 | .041 | -166.72 | -14.02 | MHS < FINS |
|  |  |  |  |  |  |  | 173.27 | 54.03 | .004 | 60.47 | 293.68 | MHS < FHS |
| Absolute | 207.14 ± 18.73 | 357.55 ± 25.74 | 334.13 ± 68.68 | 418.54 ± 35.03 | 39.42 | <.001 | 211.40 | 40.80 | <.001 | 134.10 | 293.78 | MINS < FHS |
| SO Power |  |  |  |  |  |  | 150.41 | 33.46 | <.001 | 80.40 | 216.07 | MINS < FINS |
|  |  |  |  |  |  |  | 126.99 | 72.35 | .156 | 8.47 | 271.17 | MINS < MHS |

Note. Values are reported as estimated marginals means (age controlled) ± standard error with bias-corrected confidence intervals. Only significant post-hoc analyses are presented. *p* = adjusted p; Results in bold are FDR-corrected comparisons p > .05 but became significant after bias-corrected confidence intervals. Sigma power (11.25-16Hz), SWA (0.25-4Hz), SO power (0.25-1.25Hz). NREM2 + NREM3
